# Thermal tolerance traits of individual corals are widely distributed across the Great Barrier Reef

**DOI:** 10.1101/2024.01.28.576773

**Authors:** Hugo Denis, Line K Bay, Véonique J.L Mocellin, Melissa S Naugle, Gaël Lecellier, Steven W Purcell, Véronique Berteaux-Lecellier, Emily J Howells

## Abstract

Adaptation of reef-building corals to global warming depends upon standing heritable variation in tolerance traits upon which selection can act. Yet limited knowledge exists on heat tolerance variation among conspecific individuals separated by meters to hundreds of kilometers. Here, we performed standardized acute heat stress assays to quantify the thermal tolerance traits of 768 colonies of *Acropora spathulata* from 14 reefs spanning 1060 km (9.5° latitude) of the Great Barrier Reef. Thermal thresholds for photochemical efficiency and chlorophyll retention varied considerably among individual colonies both among reefs (∼6 °C) and within reefs (∼3 °C). Although tolerance rankings of colonies varied between traits, the most heat tolerant corals (i.e. top 25% of each trait) were found at virtually all reefs, indicating widespread phenotypic variation. Reef-scale environmental predictors explained 12–62% of trait variation. Corals exposed to high thermal averages and recent thermal stress exhibited the greatest photochemical performance, likely reflecting local adaptation and stress pre-acclimatization, and the lowest chlorophyll retention suggesting stress pre-sensitization. Importantly, heat tolerance relative to local summer temperatures was the greatest on southern reefs suggestive of higher adaptive potential. These results can be used to identify naturally tolerant coral populations and individuals for conservation and restoration applications.

## Introduction

Global warming modifies species thermal environments^1^ and directly impacts organism fitness^2^. The persistence of species and the ecosystem services they provide therefore depends on their response to rising and extreme temperatures. Yet, considerable intraspecific variation in thermal tolerance exists in marine ecosystems^3^ as some species have colonized large geographical areas spanning contrasted thermal environments^4^. Individuals can adjust to local temperatures within their lifespan through physiological acclimatization and selection for particular phenotypes within populations can lead to genetic adaptation across generations^5^. Documenting phenotypic differences along thermal gradients can thus inform conservation and restoration efforts^6^ by improving predictions of species adaptive potential^3^, climate refugia^7^ or extinctions^8^.

Reef-building (scleractinian) corals are among the organisms most at risk from global warming as they occupy narrow thermal niches in environments where current summer temperatures often approach and increasingly exceed their upper thermal limits^9^. The energetic requirements of corals rely on a finely-tuned symbiotic relationship with photosynthetic dinoflagellates (family Symbiodiniaceae)^10^ that can be severely impaired under heat stress^11^, triggering a loss of symbiont cells and/or their photosynthetic pigments known as coral bleaching^12,13^. Increasingly frequent mass bleaching events have been observed over the past 4 decades^14,15^ and are predicted to have impacted > 70% of the world’s coral reefs^16^, leading to widespread coral mortality^17,18^. Understanding the extent of variation in coral heat tolerance across phylogeny, space, and time is therefore crucial to predict the future of these ecosystems^19–22^.

Intraspecific variation in coral heat tolerance exists at local scales^23–29^ and is an indicator of adaptive capacity and metapopulation persistence probability^30,31^. Populations with low intraspecific variation may suffer near extirpation at local scales^18,32,33^. On the contrary, high intraspecific variation in physiological traits can buffer populations against the loss of genetic and functional diversity^25^. Coral thermal tolerance traits are shaped by thermal history^34–36^ and other environmental conditions (e.g., irradiance^37^, hydrodynamic regimes^38^, nutrients^39^ and water oxygen content^40^). Coral colonies can adjust to these variable conditions through phenotypic plasticity^41,42^, heritable genetic effects^28,43,44^, differences in symbiont communities^27,45^ or a combination of these three factors^46,47^. Yet, further research is required to understand the relative importance of these drivers across environments, species and regions.

Quantifying intraspecific variation in thermal thresholds during natural marine heatwaves is challenged by differences in heat stress magnitude and duration over space and time and the technical constraints of surveying large spatial areas during a single heatwave event. Consequently, controlled experiments have been widely used to characterize intraspecific variation in thermal tolerance among reefs^48–52^, habitats^29,47,53^, and individuals^54,55^. Recently, acute heat stress assays (< 48 h) have been developed to measure individual thermal thresholds in a standardized, high-throughput and cost-effective manner^48–51,53,56–60^, yielding consistent results across weeks^61^, seasons^56^ and with long-term heat stress^53,57^, but see^60^). In these experiments, coral heat tolerance has been mostly inferred from photosynthetic efficiency (*F_v_/F_m_*), a common proxy of bleaching onset^62^. However, the role of symbiont photochemical damage in bleaching initiation has been increasingly challenged^63,64^ and can be offset by host protective mechanisms^12,13,65^ stressing the importance of relating *F_v_/F_m_*measurements to subsequent coral bleaching responses (i.e., the loss of symbiont cells and/or chlorophyll). To date, acute heat stress assays have investigated the response from a few colonies at many reefs or many colonies at a single reef, but have not yet evaluated intraspecific variation across multiple spatial scales.

To address the aforementioned knowledge gaps, we used acute heat stress assays to measure photochemical efficiency and bleaching traits in 768 colonies of *Acropora spathulata* from 14 reefs spanning 1,060 km (∼9.5 degrees of latitude) of the Great Barrier Reef (GBR; Figure 1a). *A. spathulata* is an abundant corymbose species found on reef flats and upper slopes that builds ecologically important structural complexity on the GBR (Figure 1d) ^66^. We identify reefs where heat tolerant individuals are most likely to be found and use multiple environmental datasets (i.e., satellite observations, numerical models, in-situ loggers) to determine environmental predictors of tolerance traits. The large sample size of this study provides GBR-wide information to inform conventional management via spatial and temporal protection and novel interventions, including seeding heat-tolerant corals onto reefs.

**Figure 1.**
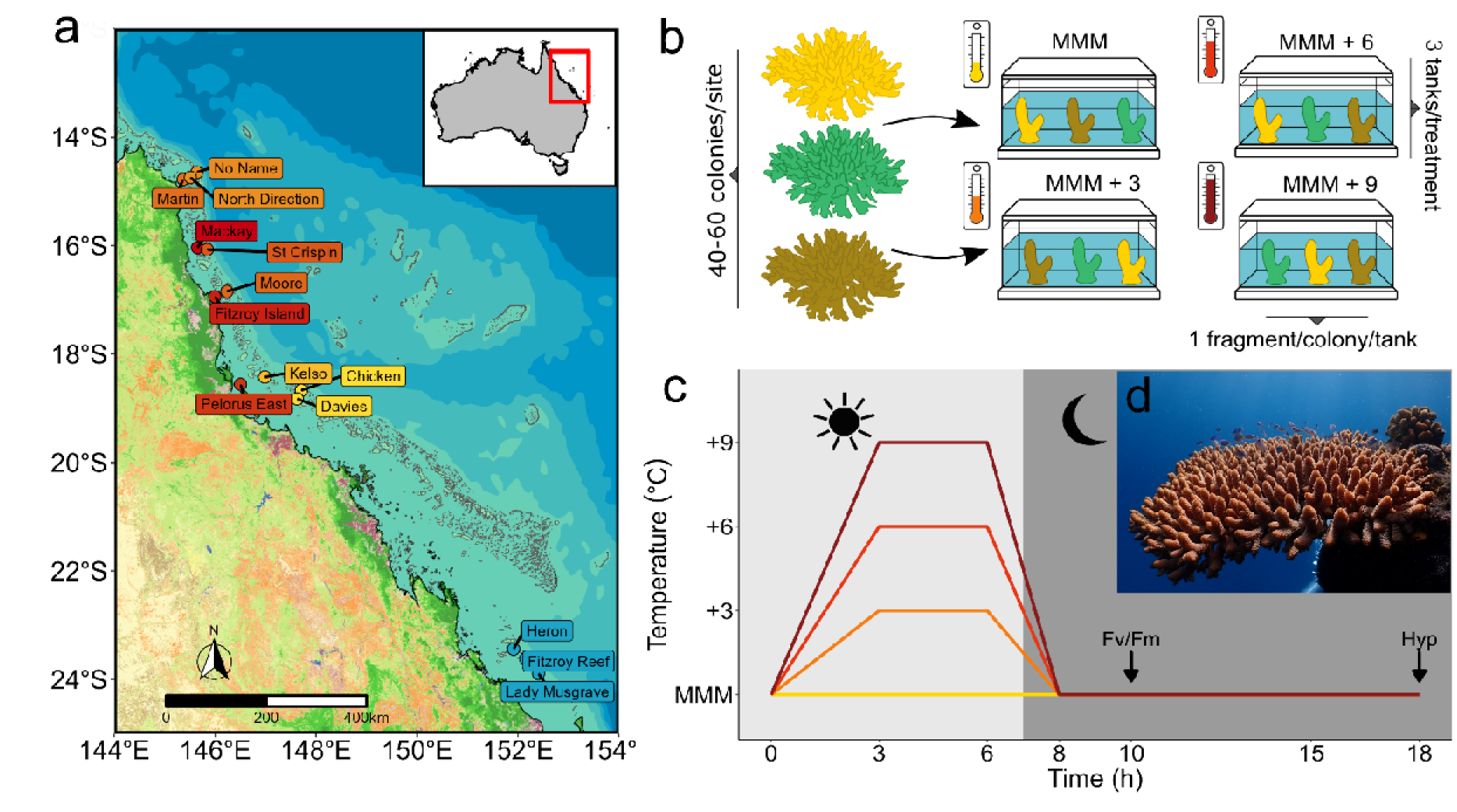
Location of sample sites and acute heat stress design to quantify variation in the heat tolerance of the reef-building coral *Acropora spathulata* on the Great Barrier Reef. **a.** Map of the 14 reefs sampled across a latitudinal and thermal gradient. Isobaths of 0, 200, 1,000 and 2,000 m were obtained from the naturalearth 10 m resolution dataset and are represented in shades of blue. Coral reef locations were obtained from the UNEP database^68^. Sites labels are colored according to their maximum monthly mean (MMM) (dark red: > 28.5 °C, blue: < 27.5 °C, Table S1). **b.** A total of 768 coral colonies were measured in standardized acute heat stress experiments. 12 nubbins per colony were randomly divided among 3 tanks for each of 4 temperature treatments relative to the local MMM of each reef. **c.** Standardized target temperature profiles to elicit increasing levels of heat stress, following Evensen *et al.* ^57^. Arrows indicate the timing of physiological measurements of the maximum quantum yield of photosystem II (*F_v_/F_m_*) and hyperspectral imagery (Hyp). **d.** Representative image of *A. spathulata* at Fitzroy Reef (© Mila Grinblat 2022).

## Results

Acute heat stress assays were performed using standardized temperature treatments to elicit increasing levels of heat stress of 0 °C (control), 3 °C, 6 °C, and 9 °C above the local Maximum Monthly Mean (MMM) ^67^ (Figure 1b). Following exposure of replicate fragments of *A. spathulata* colonies to each treatment, two phenotypic traits were measured: the maximum photochemical efficiency of photosystem II (*F_v_/F_m_*) and hyperspectral imagery estimate of chlorophyll content (normalized difference vegetation index; NDVI; Figure 1c). The decline of each colony trait was modeled over temperature treatments using dose response curves and was used to calculate two colony-level metrics of heat tolerance: the temperature that resulted in a 50% decline in response (i.e., the median effective dose, or ED50) which represents an absolute thermal threshold, and the performance under extreme heat (+9 °C), which represents how coral traits decline relative to their local reef MMM.

The majority of *A. spathulata* colonies on the GBR showed minimal decline in photochemical efficiency (*F_v_/*F_m_) and chlorophyll content (NDVI) at +3 °C (≤ 0.5%) and +6 °C (≤ 2.5%) above their local MMM. However, under the more extreme temperature of +9 °C, these traits declined by 2–80% (*F_v_/*F_m_) and 10–98% (NDVI), driving differences in thermal threshold (ED50) and performance retention metrics among reefs and individuals (Figure 2).

**Figure 2.**
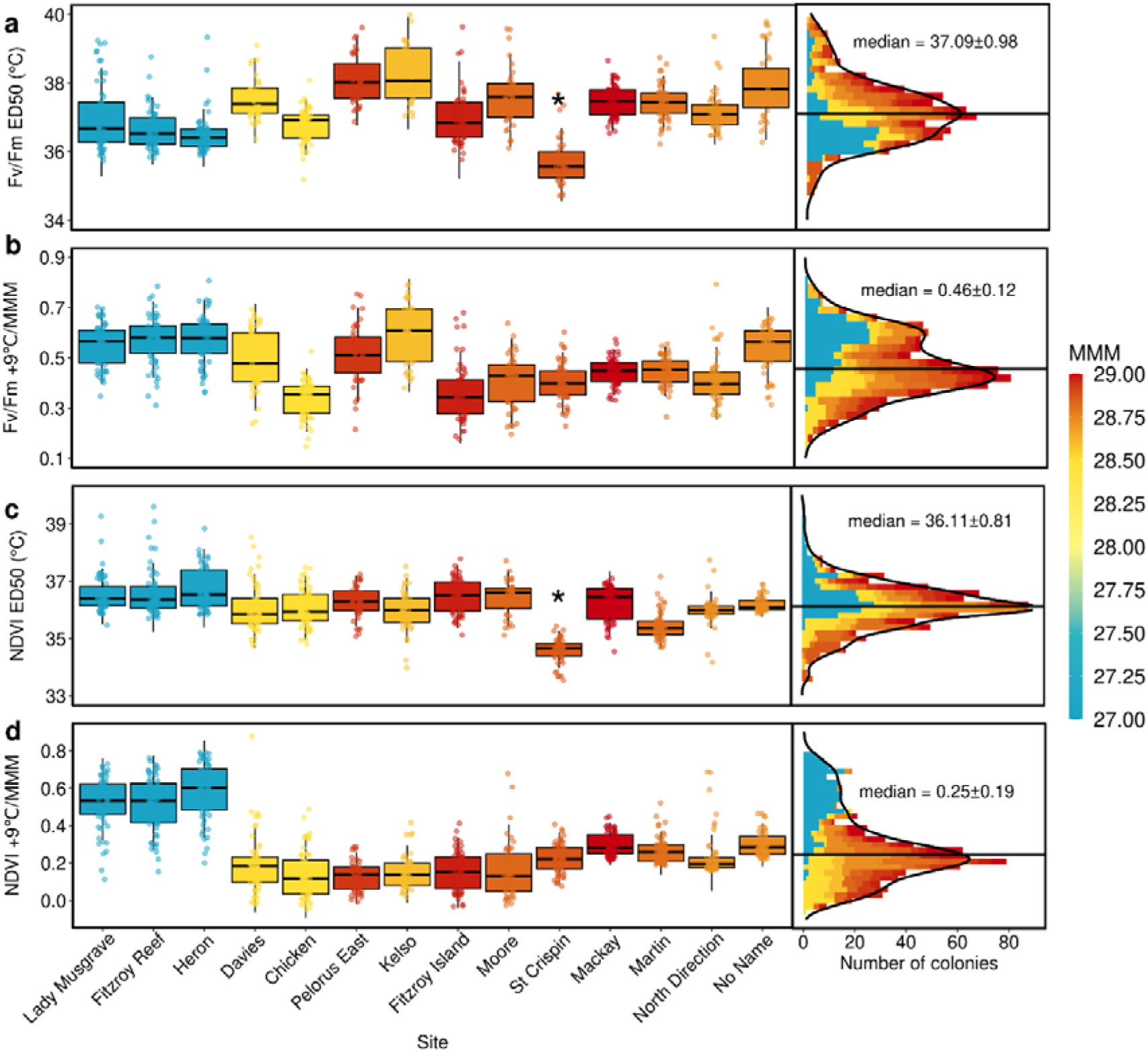
Variation in *Acropora spathulata* thermal tolerance among 14 reefs on the Great Barrier Reef (GBR) under acute experimental heat stress. The decline in maximum quantum yield (*F_v_/F_m_*) and total chlorophyll content (NDVI) was used to measure ED50 temperatures (a, c) and performance retention under extreme heat (i.e., +9 °C/MMM; b, d). Left panels display boxplots of the median (solid line) and first and third quartiles (box) for each reef colored by their MMM. Right panels show the distribution of values, with the GBR-wide median and standard deviation. Reefs are ordered (left to right) by increasing latitude. **\*** Values for St Crispin Reef are negatively biased due to experimental treatments not reaching target temperatures.

### Reef-level variation in coral heat tolerance

The GBR-wide medians of colony thermal thresholds were 37.1 ± 1.0 °C for *F_v_/F_m_* ED50 (median ± SD) and 36.1 ± 0.8 °C for NDVI ED50, with reef accounting for 34% and 25% of their respective total variation (ANOVA ω^2^ effect sizes, Table S5b). Mean reef-level *F_v_/F_m_*ED50s differed by 1.8 °C and generally decreased with latitude (Figure 2a). For example, *F_v_/F_m_* ED50 thresholds in the southern GBR (Heron Island and Fitzroy Reef; 36.5–36.6 °C) were 0.8 °C lower than all reefs in the central and northern GBR (36.8–38.3 °C; *p.adj* < .0001; Table S5d). However, several reefs had higher (Kelso Reef, Pelorus Island, No Name Reef) or lower (Fitzroy Island, North Direction Island) ED50s than nearby reefs at similar latitudes (*p.adj* < .05). Conversely, NDVI ED50 thresholds were less variable among reefs (by 1.3 °C) and were lower at Martin Reef in the northern GBR than all other reefs (*p.adj* < .01; Figure 2c). At St Crispin Reef, both *F_v_/F_m_* ED50 and NDVI ED50 were 0.8–2.6 °C lower than all other reefs but are likely to be negatively biased due to experimental treatments not reaching target temperatures (Table S2).

GBR-wide performance retention (performance under +9 °C relative to local conditions) was 46 ± 12% for *F_v_/F_m_* and 25 ± 19% for NDVI with reef accounting for 44% and 59% of their respective total variation (ω^2^ effect sizes, Table S5b). Performance retention was 2–41% (*F_v_/F_m_)* and 60–65% (NDVI) higher at southern reefs than the rest of the GBR (Figure 2b and Figure 2d), thus following an opposite trend to *F_v_/F_m_* ED50.

### Colony-level variation in coral heat tolerance

At the colony-level, thermal thresholds varied by up to 6°C, approximately 3 times the reef-level variation (1.8 °C). Within reefs, there was a 3.3 °C and 3.0 °C average range in colony-level ED50s for *F_v_/F_m_* and NDVI, respectively (Figure S1). The magnitude of this variation differed among reefs (*p* < 0.001, Levene’s test; Table S5b) but no specific reef exhibited higher or lower variation for both traits. Colony-level variation in performance retention under +9 °C was also higher than reef-level variation (by 48% for *F_v_/F_m_* and by 17% for NDVI). Although colony-level ED50s were prone to uncertainty (median 95% CI range = 2.51 ± 3.34 °C and 2.99 ± 5.56 °C, Supplementary material and methods), congruent patterns between ED50 and performance retention support high variation in acute heat tolerance among and within reefs of the GBR.

Importantly, heat tolerant colonies were widely distributed across the GBR. For example, colonies that were ranked within the top 25% of *F_v_/F_m_* ED50 and NDVI ED50 occurred at virtually all reefs (Figure 3a, b). However, the most tolerant colonies were generally more abundant at central and northern reefs for *F_v_/F_m_* and southern reefs for NDVI. Additionally, several reefs exhibited a high proportion of tolerant individuals for both traits (e.g., Mackay Reef, Moore Reef, and Pelorus Island; Figure 3).

**Figure 3.**
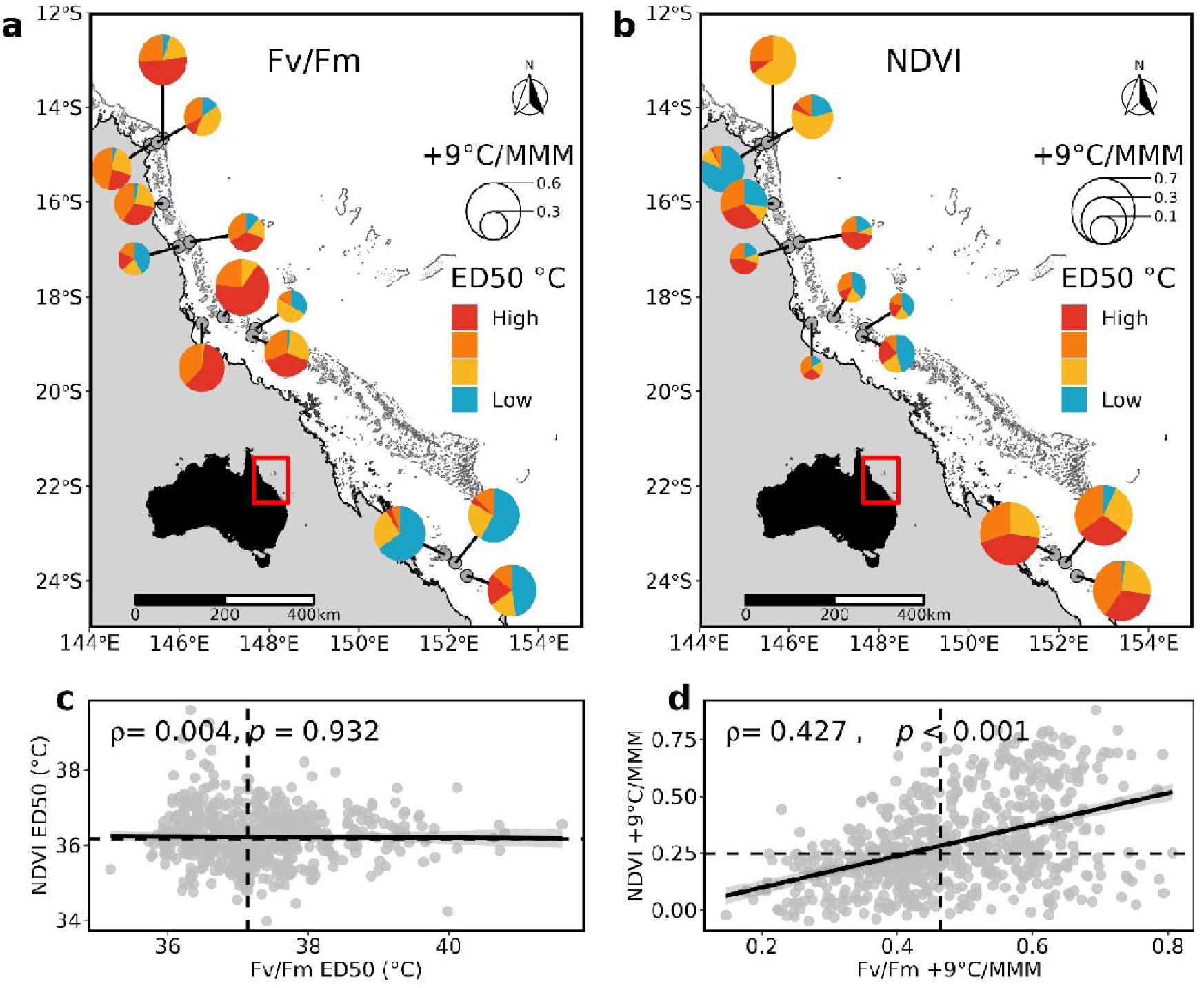
Spatial distribution of *Acropora spathulata* acute heat tolerance on the Great Barrier Reef measured by **a.** maximum quantum yield (*F_v_/F_m_*) and **b.** chlorophyll content (NDVI). Pie charts at each reef represent the proportion of individual colony ED50s falling between each quartile of the global distribution (blue: ED50 < Q1, yellow: Q1 < ED50 < Q2, orange: Q2 < ED50 < Q3, red: Q3 <ED50). Pie charts radii are proportional to average retained performance under extreme heat (i.e., +9 °C/MMM), normalized by the global range for plotting purposes. Correlation between **c.** *F_v_/F_m_* and NDVI ED50s and **d**. retained performance under extreme heat (+9 °C/MMM). Each point represents a distinct colony (n = 652 and 730 for c and d, respectively) and Spearman’s-ρ and *p-*values are indicated.

### Response to acute heat stress differs between traits

The ranking of reef- and colony-level responses under heat stress differed between phenotypic traits. Overall, there was no GBR-wide correlation between *F_v_/F_m_* and NDVI ED50s (*ρ* = 0.00, Figure 3c), but a moderate positive correlation between their performance retention at +9 °C (*ρ* = 0.43, Figure 3d). At the reef-level, there was a moderate positive correlation between *F_v_/F_m_* ED50 and NDVI ED50 at Chicken and No Name (*ρ* > 0.3) and a weak positive correlation at Davies, Heron, Mackay, Martin and Moore (*ρ* = 0.1–0.3, Table S6). Similarly, there was a weak to moderate positive correlation between the performance retention of *F_v_/F_m_* and NDVI at all reefs (*ρ* = 0.1–0.3) except Chicken (*ρ* = 0.02), North Direction (*ρ* = 0.84) and Pelorus Island (*ρ* = −0.22; Table S6).

### Environmental drivers of acute heat tolerance metrics

To examine environmental conditions experienced by *A. spathulata* across the GBR, we retrieved 10 environmental variables characterized by 24 quantitative predictors computed from *in-situ* loggers, numerical models, and satellite observations (Table S7). A Principal Component Analysis on environmental predictors divided the 768 colonies into three major clusters (Figure S2). The first component (46.1% explained variance) separated cooler southern reefs from warmer central and northern sites and was mainly driven by MMM (1.7 °C range) and Degree Heating Weeks (DHW) at the time of collection. The second component (15.1% explained variance) separated northern and central sites along an inshore-offshore gradient that was mainly driven by turbidity, sea surface current velocities, and temperature variation (annual range and the rate of change from spring to summer). Within clusters, colonies grouped by reefs as most predictors were retrieved at the site level and were minimally separated by their water depth or pigmentation. Pigmentation scores prior to collection differed between individuals and sites but had no clear association with DHW at the time of collection (Figure S3).

Environmental drivers of heat tolerance traits (ED50 and performance retention in *F_v_/F_m_* and NDVI) were evaluated using random forest ensemble learning and ridge regression on 12 low to moderately correlated predictors (pairwise absolute Pearson correlation = 0.01–0.69). Together, these predictors explained a moderate to large proportion of variation in heat stress responses of *A. spathulata* across the GBR (*R*^2^ = 0.12–0.62). The predictive accuracy of random forest (RF) and ridge regression (RR) was stronger when heat tolerance was expressed for both *F_v_/F_m_* and NDVI as performance retention (*R*^2^ = 0.27–0.62) than ED50 thresholds (*R*^2^ = 0.12–0.31). Top environmental predictors slightly differed among traits and metrics but variation in heat tolerance was primarily associated with site-level thermal history (Figure 4). MMM (Figure 4b) and DHW at time of collection (Figure 4d) were predominant metrics associated with response to acute heat stress (i.e., top RF predictors for 3 of the 4 heat tolerance metrics, Table S10a) while being strongly correlated to each other (*R* = 0.78). The direction of associations differed between traits with MMM and DHW at the time of collection being positively associated with *F_v_/F_m_* ED50 (β = 0.06–0.07), but negatively with performance retention in *F_v_/F_m_* and NDVI and NDVI ED50 (β = −0.03–-0.06; Table S10b). Secondary thermal history associations occurred between the annual range in temperature and both performance retention metrics (*F_v_/F_m_* β = −0.03, NDVI β =-0.04; Figure 4c), and between the frequency of DHW > 4 and both ED50 thresholds (*F_v_/F_m_* ED50 β = 0.04, NDVI ED50 β = −0.06; Figure 4e). Particularly, the frequency of DHW > 4 in the year before collection showed a strong negative association with NDVI ED50 (β = −0.060, top predictor in RF), but no significant association with any other trait metric.

**Figure 4.**
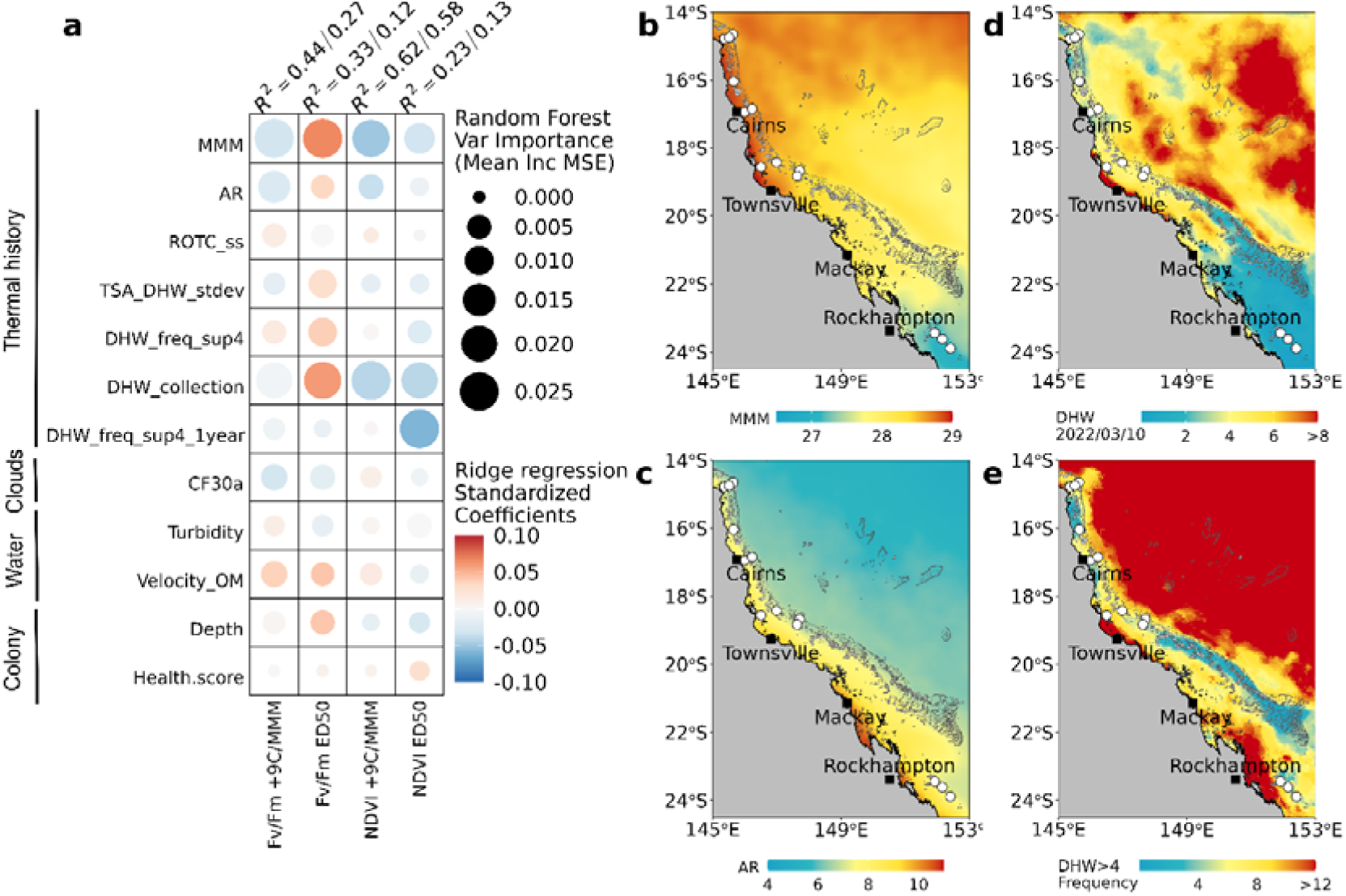
Environmental drivers of acute heat tolerance of *Acropora spathulata* on the Great Barrier Reef. Heat tolerance was measured as the maximum quantum yield of photosystem II (*F_v_/F_m_*) and chlorophyll content (NDVI) expressed as ED50 temperature thresholds and performance retention under extreme heat (+9 °C). Relationships between environmental predictors (defined in Table S7) and heat tolerance metrics were assessed using ridge regression and random forest models. **a.** Heatmap of associations evaluated with random forest variable importance (10-fold stratified cross validation, circle size) and ridge regression standardized coefficients (multiple bootstrap resampling *n* = 100, circle color). The variance explained by each model (random forest/ridge *R*^2^) is shown above each tolerance trait metric. **b-e**. Thermal history maps of (**b**) maximum monthly mean temperature as defined by NOOA^67^ (**c**) annual temperature range (2014–2022), (**d**) peak degree heating weeks during sampling and (**e**) annual frequency of degree heating weeks above 4 since 1985. White dots indicate reef sites sampled in this study.

To a lesser extent, the heat tolerance of *A. spathulata* was associated with water flow, shading, and depth. *F_v_/F_m_* tolerance metrics were generally higher at reefs with high sea surface current velocity (*F_v_/F_m_*ED50 β = 0.042, +9/MMM β = 0.037) and lower at sites with positive cloud anomalies in the 30 days prior to collection (*F_v_/F_m_* ED50 β = −0.017, +9/MMM β = −0.031; Figure 4a). Turbidity showed little to no relationship with heat tolerance metrics. At the colony level, increasing water depth was positively related to *F_v_/F_m_* metrics (*F_v_/F_m_* ED50: β = 0.042, +9/MMM: β = 0.004), but negatively related to NDVI metrics (NDVI ED50: β = − 0.028, +9/MMM: β = −0.015). In addition, colonies with higher levels of pigmentation at the time of collection tended to have higher NDVI ED50 thresholds (β = 0.028), but there was no relationship with other tolerance metrics.

## Discussion

The large geographic scope of this study (14 reefs across 9.5 degrees of latitude) and sample size (*n* = 768) revealed extensive intraspecific variation in the heat tolerance of reef-building corals across the Great Barrier Reef (GBR). This resulted in more than 6 °C variation in colony thermal thresholds where tolerant colonies — based on multiple traits and metrics — were widely distributed among and within reefs. Environmental predictors of heat- tolerance variation support adaptation and/or acclimatization of populations to local conditions while unexplained differences among colonies may be due to host and/or symbiont adaptive genetic variation within reefs.

### Acute heat tolerance metrics linked to historical and recent environmental conditions

All heat tolerance metrics were most strongly related to thermal history supporting adaptation of corals to their local thermal regimes^35,36,69,70^. These results also corroborate ecological experiments that found 4 °C difference in *A. spathulata* larvae survival thresholds between Lizard Island (14.7 °S) and One Tree Island (23.5 °S) i.e., the same latitudinal range as our study^71^. *F_v_/F_m_* ED50 thresholds increased at reefs exposed to higher thermal averages (MMM), annual thermal variation (AR) and exposure to marine heat waves (frequency of DHW > 4 and standard deviation of DHW). DHW at the time of sampling was also strongly positively associated with *F_v_/F_m_* ED50, suggesting that acclimatization of the holobiont to higher temperatures possibly through upregulation of photoprotective mechanisms can increase the heat tolerance trait *F_v_/F_m_* ED50 as reported by Cunning *et al.* ^56^. Conversely, there were weaker unexpected negative associations between all thermal history metrics and NDVI ED50. This may reflect accumulated heat stress leading to early health impairment of the holobiont and lower chlorophyll retention, which is supported by the strong negative relationship of NDVI ED50 with frequency of DHW >4 in the 12 months prior to sampling.

This is similar to *F_v_/F_m_* ED50 findings of Marzonie *et al.* ^49^ in the Coral Sea, suggesting that some reef populations may be impacted over long time periods after heat exposure. Differences in the association with thermal history among traits could thus occur from a decoupling between host and symbiont baseline conditions and/or stress responses (discussed below)^63,64,65^. Nevertheless, several reefs in our study (No Name, North Direction, Mackay) exhibited a high proportion of tolerant individuals for both *F_v_/F_m_* and NDVI ED50, showing that some colonies and populations may perform better under heat stress across multiple traits^72^. These populations will be more likely to play a key role in the GBR resilience in a warming climate and thus their mapping in space and time could provide valuable information for spatial planning of Australia’s marine parks into the future.

The heat tolerance metrics of *A. spathulata* measured here were further associated with predictors that influence water movement, light, and within-reef temperature. Strong water movement can delay photochemical damage under thermal stress through coral surface cooling, increases in respiration and metabolic transfers and removal of toxins^38,73^ and the positive association between surface current velocity and *F_v_/F_m_* metrics suggests a carryover of these effects to our experiments. Solar irradiance is also a factor known to mediate bleaching responses under high seawater temperatures^74^ and *F_v_/F_m_* ED50 thresholds were indeed reduced at reefs with a high cloud cover prior to the experiments. This may indicate that symbionts acclimated to lower light levels^75^ experienced greater irradiance-induced photosynthetic damage in the experimental system^76^. Overall, these results demonstrate that empirically derived thermal tolerance metrics are not only shaped by long-term adaptation but also acclimatization to recent conditions.

### Standing within-reef variation in heat tolerance supports adaptive potential to climate change

There was greater variation in the acute heat tolerance of *A. spathulata* colonies within reefs than among reef-level averages (ED50 ∼3.3 vs 1.8 °C and ∼3.0 vs 1.6 °C for *F_v_/F_m_* and NDVI respectively), despite most colonies being sampled from a narrow depth range at a single site less than 600 m^2^. Importantly, we found that heat-tolerant corals (defined as the top 25 percentile from the GBR ED50 distributions) occurred at nearly every reef, including the cooler southern GBR. These results align with previous results on *A. cervicornis* in the Florida Keys^56^, *A. hyacinthus* in Palau^51^ and *A. hyacinthus* in the GBR^77^. Within-reef variation remained mostly unexplained by colony-level predictors (depth and pigmentation) and could be elucidated by incorporating genomic data not yet available here. Individual differences in heat tolerance may be underpinned by genomic variation within the host^23,28^, symbionts^45,78^ and/or their interactions^55^. For example, incorporating the proportion of *Durusdinium* cells and host polygenic scores based on putative heat-adaptive loci increased the predictive accuracy of GBR *A. millepora* bleaching models by ∼22%^79^. In addition, deciphering the independent role of environmental drivers requires a better characterization across microhabitats^80^ and depth^81^ as well as optimization of sampling strategies across environmental gradients to reduce the collinearity of predictors^82^. The large standing variation in thermal tolerance at the reef scale reported in this study has important implications for the conservation of this species on the GBR. It may notably support *A. spathulata* adaptation to climate change through selection of pre-existing heat tolerance alleles^20^. Furthermore, rather than translocating individuals from warmer reefs (e.g., Howells *et al.* ^69^) restoration projects can target local heat tolerant individuals when their phenotypes are known or by sampling a diversity of genets.

### Coral populations from the Southern GBR may live further from their upper thermal limit

Acute thermal thresholds (ED50) for *A. spathulata* populations on the GBR were 9.5–9.6 °C above their local MMM on southern reefs compared to 7.5–8.8 °C on central and northern reefs. This is consistent with higher performance retention under extreme heat on southern reefs and latitudinal differences in the thermal thresholds of *A. spathulata* larvae^83^.

Consequently, under comparable levels of thermal stress, southern populations are expected to be less susceptible to bleaching than central and northern populations. In line with this expectation, southern GBR reefs have experienced less frequent and intense bleaching events than central and northern reefs^15^ with 90% less bleaching-related mortality during the 1985–2012 period^17^. The lower frequency and magnitude of bleaching events on southern reefs might also allow more time for adaptation to occur through the migration of heat-tolerance associated alleles from warmer northern reefs^20^.

Conversely, the low cumulative heat stress experienced by those regions may result in coral assemblages that are naïve to high heat stress^15^. As such, mild but highly prevalent coral bleaching occurred in the southern GBR during 2016–2017 (21% of colonies; Kennedy et al. ^84^) and 2020 (86% of colonies; Page et al. ^85^; 48% of colonies; Nolan et al. ^86^) and occurred under lower cumulative heat stress (DHW) than in central and northern GBR^15^. Further investigations are thus required to understand how recurrent warming disturbances and gene flow with northern regions will shape the evolution of these cooler reefs.

### Divergent effects of acute heat stress on thermal tolerance traits

Here we found a moderate correlation between *F_v_/F_m_* and NDVI performance retention under extreme heat. This supports the use of photochemical apparatus integrity as an early proxy of coral bleaching^53,57,62^ and the importance of symbiont tolerance in bleaching response. For instance, some symbiont species or genotypes may produce more photoprotective pigments or antioxidant compounds, alleviating oxidative stress for the host ^12,13^. Conversely, we found no correlation between ED50s of *F_v_/F_m_* and NDVI traits. The only two other studies using *F_v_/F_m_* and chlorophyll-related ED50 thresholds found congruent reef-level variation between the two traits for some species (e.g., Pocillopora verrucosa^48^ and *Acropora hyacinthus*^77^) but not others (e.g., Porites lobata^48^). In both studies, visual chlorophyll scores showed higher noise and lower differences in reef ED50s which aligns with *A. spathulata* pairwise reef differences in NDVI ED50s (0.39 °C) being almost half that of *F_v_/F_m_* ED50 (0.67 °C). This may also be due to differences in the timing of measurement between traits as their decline occurs at different rates^63^. In *A. tenuis*, *F_v_/F_m_* values were found to be stable 0–24 hours after the end of an acute heat stress, while chlorophyll decreased up to 24 hours after the end of temperature ramp-down^87^. Therefore delaying the timing of NDVI measurements in our study may have revealed higher divergence in chlorophyll retention between reefs. Finally, algal oxidative stress and photosynthetic damage can be alleviated by antioxidant mechanisms^13^ or immune response from the host^65^ and thus occur without a decrease in chlorophyll content or symbiont density^88^.

Divergent responses between traits highlight the importance of defining heat tolerance and selecting phenotypic traits accordingly. Chlorophyll content may be an appropriate proxy for holobiont bleaching and mortality as its decline during an acute heat stress can persist and reflect mortality in the next month^61^. However, photosynthetic efficiency may be a better proxy to detect sub-lethal effects as corals can experience physiological stress of reduced tissue biomass and symbiont loss long before changes in pigmentation are visually detectable^63^. Finally, which phenotypic trait under acute experimental stress would best predict heat tolerance in natural conditions remains unclear and could vary with marine heatwave magnitude and duration as well as the biological level at which they are assessed^60^. Our results demonstrate that no single trait or metric can fully capture the complexity of the coral holobiont heat stress response, and highlight the importance of measuring multiple traits whenever achievable, including relevant host traits (e.g., antioxidant capacity).

## Conclusion

Using standardized acute heat stress assays, we found that heat tolerant corals were widely distributed across the GBR, including cooler, southern reefs. This suggests potential for adaptive responses to climate change, if heat tolerance traits have a heritable basis^31,44^. Rankings of coral heat tolerance can guide human interventions and their interpretation and application may depend on the specific traits and metrics measured. Corals with high absolute thermal tolerance can help to decipher heat-stress protective mechanisms^23,89^ and be used as material for assisted gene flow through translocation ^59,90^ and selective breeding^43^. On the contrary, corals with high thermal tolerance relative to local conditions may be better targets for local propagation and breeding, decreasing the risk of carrying pathogens and phenotype-environment mismatches^91^. Nonetheless, natural adaptive processes and human interventions must be accompanied by reduced anthropogenic emissions through ambitious national and international commitments to secure the persistence of species and populations across the entire GBR.

## Methods

### Study sites, experimental design and setup

#### Study sites and sample collection

We quantified thermal tolerance traits of 768 colonies of *A. spathulata* across 14 reefs of the GBR (Figure 1a, Table S1). At each reef, 40–60 colonies were sampled on SCUBA from one or two sites (covering 50–4200 m^2^) between 28 February and 26 March 2022 (GBRMPA permit G21/45166.1) over a single or two consecutive days. For each colony, 12 nubbins (∼8 cm) were collected for standardized acute heat stress experiments and *in-situ* metadata was recorded. This included colony-level GPS coordinates linked to *in-situ* photographic records^92^, time-corrected depth (−0.54 ± 0.76 m relative to the lowest astronomical tide) and visual pigmentation (Coral Watch Health Chart scores at 0.5 intervals)^93^. Colonies showing signs of disease were rarely encountered but intentionally excluded to avoid physiological bias unrelated to temperature effects.

#### Acute heat stress assay design

A portable automated experimental system (“Seasim-in-a-box”, National Sea Simulator, XXX; Figure S4) was modified from Marzonie *et al.* ^49^ to conduct acute heat stress assays onboard a research vessel the day following collection (Supplementary materials and methods). The experimental design consisted of three replicate tanks for each of four temperature treatments of 0 °C (control), 3 °C, 6 °C, and 9 °C above the local Maximum Monthly Mean (MMM, 1985–1990+1993 climatology as defined by NOOA^67^) for each reef site. These treatments were designed to elicit increasing levels of thermal stress following Voolstra *et al.* ^53^. Immediately after collection, 12 nubbins from each coral colony were mounted on separate experimental racks with unique identifiers and held overnight at ambient seawater temperature. The following morning, one nubbin from each colony was randomly assigned to each of the 12 tanks (Figure 1b). Assays started at 11:00 am and consisted of 3 hours ramp-up from ambient temperature (within 1 °C of the MMM) to target temperature, 3 hours hold at the target temperature, 2 hours ramp down to MMM, and a final ∼12-h hold at MMM (Figure 1c). During the experiments, temperatures in individual tanks were recorded at a 1-min intervals using Hobo loggers (Onset) and closely matched their target profiles (mean_ΔT_ = 0.36 °C, Table S2) with the exception of one site (St Crispin: mean_ΔT_ = 1.61 °C).

### Phenotypic traits

#### Photochemical efficiency

The maximum quantum yield of photosystem II (*F_v_/F_m_*) of coral photosymbionts (Symbiodiniaceae) was used as an initial rapid, non-invasive measure of heat tolerance. A decline in *F_v_/F_m_* reflects early physiological impairment in corals and has been reported to respond consistently between acute- and long-term heat exposures^53,57^; but see Klepac *et al.* ^60^. Measurements were taken 2 hours after the end of temperature ramp-down (> 1-hour dark adaptation) using an Imaging PAM chlorophyll fluorometer (IMAG-K7, Walz Germany) with the following settings: Measuring Light = 2 (freq = 1), Saturation Pulse = 7 (Int = 30s), Damp = 1, Gain = 1. For each fragment, three measurements were extracted from non-overlapping areas and underwent several quality check and filtration steps (Supplementary materials and methods). After filtration, a total of 769 colonies and 8,453 fragments were retained in further analyses.

#### Hyperspectral image-based assessment of chlorophyll content

Non-invasive estimates of chlorophyll content were used as a second metric of heat tolerance. The loss of chlorophyll underpins visual bleaching scores commonly used in acute heat stress experiments and has been shown to correspond with mortality risk^61^. Total chlorophyll content was assessed in fragments from reflectance measurements taken 11 hours after the end of the temperature ramp-down using a hyperspectral camera (Resonon, Pika XC2) with the following settings: Integration time = 39.2ms, Gain = 0, Frame rate = 22 fps. MATLAB software was used to compute the normalized difference vegetation index (NDVI, Supplementary material and methods) from reflectance measurements, where NDVI = (R_720_-R_670_)/(R_720_+R_670_) and R_x_ is the reflectance at x nm. NDVI is a proxy for chlorophyll used in a wide range of organisms including scleractinian corals^94^ and has been validated in the soft coral *Sarcophton* cf. *glaucum*^95^. We repeated this validation in *A. spathulata* using spectrophotometric determination from tissue extractions^96^ with a strong relationship between NDVI and log-transformed chlorophyll-a (*R*^2^=0.74, Supplementary material and methods, Figure S5).

### Phenotypic data analysis

The response of raw *F_v_/F_m_* and NDVI to temperature across experiments was assessed using a linear mixed effect model incorporating fixed effects of site, treatment, fragment size (estimated from pixel numbers in RGB images) and random intercepts for each colony and tank using package *lme4*^97^, including 95% bootstrap confidence intervals for fixed effects (Table S3). Tanks were first treated as a block factor in the analysis. Despite small variations in temperatures among replicate tanks, tank had a minor effect on fragment performances under heat stress (explained 1.44% of *F_v_/F_m_*random variance σ_i_ ^2^ and 0.8% of NDVI σ ^2^, linear mixed effect model, Table S3). Likewise, fragment size effect appears to be negligible for *F_v_/F_m_*(B = 1.4 × 10^-6^, df = 7473, 95% CI: −4.8 × 10^-7^–3.0 × 10^-6^) and very weak for NDVI (B = 8.5 × 10^-6^, df = 8988, 95% CI:6.14 × 10^-6^–1.0 × 10^-5^).

#### ED50 thermal thresholds

For each colony, the decrease in *F_v_/F_m_* and NDVI was modeled over temperature treatments using dose response curves in the R package *drc*^98^. Colonies with <2 fragments per treatment after previous quality control filtering (Supplementary material and methods) were excluded to ensure robust estimates of individual responses across treatments (excluded colonies: *F_v_/F_m_* = 103, NDVI = 4). All models were fitted using the *drm* function based on the mean hold temperature recorded within each tank for each experiment, with constraints set on parameters (Table S4a). The best model (Weibull type II with 3 parameters) was chosen as the one with the lowest Akaike’s information criterion score (AIC) for most reef sites using the *mselect* function (6/14 and 11/14 reefs for *F_v_/F_m_* and NDVI, respectively; Table S4b).

Because using the same model is a prerequisite to compare ED50s, we used this model for all colonies even though some curves showed a better fit to log-logistic or quadratic models.

The model equation is

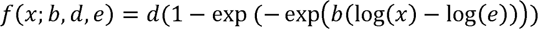

where *b* is steepness of the curve, *d* the upper asymptote (lower asymptote set to 0) and *e* the inflexion point. An example of dose response curves can be found in Figure S6. Dose response curves were filtered to remove individuals that showed poor fit to the data, notably when the decline in phenotypic traits was minor or absent up to +6 °C or +9 °C resulting in ED50s with wide confidence intervals (Supplementary material and methods). After filtration 652 and 725 colony-level ED50 estimates were retained for *F_v_/F_m_* and NDVI respectively.

Performance retention under thermal stress Performance retention under extreme heat was computed as

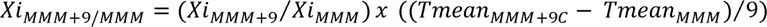

where *Xi* is the trait (*F_v_/F_m_* or NDVI) averaged across fragments of individual i in the MMM and MMM+9 treatments, respectively, and T*meanMMM_+9_C* and T*meanMMM* are the average hold temperatures for these treatments. The second term accounts for small differences between target and effective temperatures across tanks and experiments. Performance retention can be computed even for colonies that experienced minor decline in phenotypic traits under +9 °C which prevented fitting a dose-response curve. After filtration of colonies with < 2 fragments in the MMM and MMM+9C treatments, 730 and 765 colony-level performance retention estimates were calculated for *F_v_/F_m_*and NDVI respectively.

### Heat tolerance variation analysis

The variation in heat tolerance (ED50 and performance retention) among and within sites was investigated using an analysis of variance (ANOVA). Homogeneity of variance between sites was assessed using Levene’s test on residuals from groups medians (Table S5b) and Tukey’s HSD post-hoc tests for pairwise comparisons (Table S5c). Since variance was unequal across sites, we used one-way Welch’s ANOVA to compare site means and Games-Howell post-hoc tests with adjusted *p*-values at α = 0.05 for pairwise comparisons (Table S5d). Following Cunning *et al.* ^56^, we assessed within-site variation in heat tolerance by computing ED50 adjusted as the grand mean of the total distribution plus residuals from site. Finally, we investigated the similarity of colony thermal tolerance rankings between *F_v_/F_m_* and NDVI traits using Spearman’s rank correlation as assumption of normality was violated. For both ED50 and performance retention, we computed both GBR-wide and site-level correlations (St Crispin Reef excluded; Table S6).

### Environmental data acquisition and analysis

To examine environmental conditions experienced by *A. spathulata* across the GBR, we retrieved 10 environmental variables characterized by 24 predictors computed from *in-situ* loggers, numerical models, and satellite observations (Table S7). Reef thermal history was characterized using the NOAA Coral Watch v3.1 sea surface temperature (SST) satellite product (1985–present; 5km)^99^ for thermal predictors based on historical climatology (MMM, DHW) as it has the largest temporal coverage. For contemporary thermal predictors (e.g., annual average and thermal range, 2014–present) we used the eReefs GBR1 Hydro model^100^ as it delivers temperature predictions at a finer spatial resolution (∼1 km, hourly) and higher accuracy for the last decade (ME = 0.4–0.9 °C)^101^. Temperatures from both sources were validated using *in-situ* records (Supplementary material and methods).

Global averages (2010–2019) of environmental variables that may play a secondary role in heat tolerance (e.g., chlorophyll, turbidity, current velocity) were retrieved from eReefs GBR4 model (hourly; 4 km)^102^. In addition, we included the average cloud fraction (% of pixel masked by clouds) and cloud fraction anomaly for the 30 days prior to collection^37^ retrieved from NASA-MODIS 1°, monthly database^103^.

Our environmental dataset also included variables measured at the individual colony level. Depths obtained from dive computers were adjusted to tide level at the time of collection (standardized to Lowest Astronomical Tide, Figure S7). Predictors derived from eReefs were computed at each colony depth through vertical interpolation from the model z-levels. As the GBR experienced a marine heatwave during the sampling campaign (0.47–4.67 DHW), we included pigmentation levels prior to collection as an explanatory variable to account for heat stress experienced in the weeks preceding the experiment (Figure S3).

Principal Component Analysis (PCA) was performed on environmental predictors for the 768 colonies to visualize the distribution of sites and colonies along environmental gradients.

### Phenotype by environment analysis

The influence of environmental predictors on heat tolerance traits (ED50 and performance retention in *F_v_/F_m_* and NDVI) was evaluated using random forest ensemble learning and ridge regression on a set of 12 low to moderately correlated predictors (pairwise absolute Pearson correlation = 0.01–0.69; Figure S8, Table S9; St Crispin Reef excluded). Because any association of heat tolerance traits with MMM and DHW at the time of collection may reflect different mechanisms (e.g., adaptation vs acclimatization), both were retained in the dataset despite their strong correlation (*R* = 0.78).

Random forest models were built using *cforest* function from *party* R package^104^. For each of the 4 heat tolerance metrics, separate random forests were grown to 1,000 trees (ntree) with 5 environmental predictors tried at each split (mtry). Predictor importance and model accuracy were assessed through repeated 10-fold stratified (across sites) cross-validation (70/30 split, Supplementary material and methods). The importance of each predictor was estimated by computing the marginal increase in out-of-bag sample MSE (Mean Inc MSE) when training the model with the predictors randomly shuffled.

Ridge regression was performed for each phenotypic trait using the *glmnet* R package^105^. The optimal tuning parameter λ of the penalty term was selected using k-fold cross-validation (λ.1se; Figure S9 and Figure S10). Model accuracy (*R*^2^) and environmental predictor coefficients were estimated using bootstrap resampling (repeated 100-fold stratified cross validation, Figure S11 and Table S10).

All statistical analysis were conducted using R v4.0.4 and figures created using package *ggplot2* and Inkscape v1.2.

## Data and code availability statement

Scripts and data used for the analyses are available online: https://github.com/hde08/Aspat-GBR-HeatTolVar/tree/main. Data is publicly archived on [Anonymous Repository].

