## Supplementary Materials for "Thermal tolerance traits of individual corals are widely distributed across the Great Barrier Reef"

### A. Material and Methods

1. **Automated experimental system**

The portable experimental system (“Seasim-in-a-box", XXX) consists of replicate 50-L acrylic tanks outfitted with insulating water jackets and supplied with 0.8 L. min^1^ unfiltered seawater directly pumped from the ocean (Iwaki MX400). Each treatment sumps, with recirculating water supply to the insulating water jackets, were brought from ambient to target temperatures by heating elements (Omega 2kW titanium). In parallel, four distinct flow-through seawater were heated by convection via their titanium coil (Wateco 56″ titanium heat exchanger) placed in the sumps, and pumped to the replicate experimental tanks using submersible pumps (Reefe RP2400LV). Tanks were equipped with independent white/blue LED lighting (300 W) to supply ~ 300 µmol photons m^-2^. s^-1^ with a diurnal light regime timed to *in-situ* conditions (11:13). Each tank was equipped with circulators (Tunze 6015), independent temperature probes (TC Direct PT-100) and PAR sensors (Skye Quantum PAR – SKL26250) to control temperature and lighting. The monitoring and control of temperature and lighting were achieved by a system consisting of a Programmable Logic Controller (Siemens Simatic 571500), a Weidmuller UR20 Remote IO Signal Input and Outputs and a Human Machine Interface (HMI TP700 Comfort panel).

#### 2. Photosynthetic efficiency measurement

For each fragment, *Fv/Fm* values were extracted from 3 manually-selected non-overlapping areas from fluorescence images in ImagingWin v2.56 software using a custom PYTHON script to semi-automate the extraction. Fluorescence images underwent posterior quality checks and a subset of fragments was removed from the dataset due to technical artifacts. In addition, filtering steps were applied to remove i) *Fv/Fm* records above 0.81 as they exhibited an unexpected dependency between *Fo* and *Fv/Fm* (Linear regression, p<2 x 10^-16^, R^2^=0.336), ii) coral fragments with *Fv/Fm* standard deviation > 0.2 among areas leading to high uncertainty on the mean *Fv/Fm*.

#### 3. Validation of chlorophyll measurements from hyperspectral imaging

Hyperspectral images were analyzed using XXX proprietary tools in MATLAB software. Each coral fragment was manually masked on RGB images as a 2D-polygon using Image Processing Toolbox and reflectance measurements individually extracted for each wavelength from pixels belonging to each fragment. We assessed the accuracy of hyperspectral imaging to estimate chlorophyll-a content in *A.spathulata* by independently measuring chlorophyll-a on a subset of individuals originating from two sites (Chicken Reef/Pelorus Island) and MMM/MMM+6 °C treatments. To this end, snap-frozen coral fragments at the end of the experiment were freeze-dried and crushed. For each fragment, an aliquot of homogenized powder was placed in 95% ethanol then sonicated in ultrasound bath (275W, 20 kHz) and centrifuged (10 000g, 5min) to release chlorophyll content into solution. Chlorophyll a was quantified from absorbance measurements taken in a microplate spectrophotometer (Bio-tek Powerwave) using Ritchie. R. J equation ^1^ and standardized to dry weight. Finally, we performed a linear regression of NDVI on untransformed and log transformed chlorophyll-a content (Supplementary Fig. 5).

4. **Filtration of dose response curves**

For a large proportion of colonies (*Fv/Fm*: 55%, NDVI: 21%) the decline in phenotypic traits was small or absent up to +6 °C above control temperatures, and inferior to 50% under +9 °C. This resulted in predicted ED50s outside the range of temperature treatments used in the experiments and wide confidence intervals (>10 °C). Furthermore, log-logistic models could not be fitted at all for some of the most tolerant colonies because there was virtually no decline in traits even under +9 °C. In order to retain naturally tolerant individuals without artificially boosting reported variation in heat tolerance, we discarded from further analysis colonies that were both detected as an outlier based on the boxplot visualization method on the total distribution (ED50<Q1-1.5*IQ or ED50>Q3+1.5*IQ) and had a confidence interval larger than 10 °C as well as colonies for which confidence interval computation failed indicative of lack of fit to the data (*Fv/Fm*: *n* = 20, NDVI: *n* = 40). We acknowledge that some of the top performers may have been overlooked using this absolute metric, and that individual estimates of ED50s are prone to uncertainty.

#### 5. Validation of satellite and model temperature predictions

Temperature time-series from both sources were compared to *in-situ* records (1-3m depth, 13/14 sites) from *aimsdata* R package ^2^. Depending on available *in-situ* records, daily time-series spanning between 1–7 years of the 2014–2022 period were aligned against eReefs and Coral Watch predictions to compute Pearson correlation coefficients (Supplementary Fig. 12 and Supplementary Table. 8). We also assessed the accuracy of eReefs at different time resolutions (hourly, daily, weekly, Supplementary Fig. 13). Overall, there was a good agreement of satellite observations and model predictions to *in-situ* measurements at a daily resolution (R>0.92, Supplementary Fig. 12-13, Supplementary Table. 8). However, subtracting low frequency variability (daily and weekly averages) from hourly and daily temperatures resulted in low/moderate to strong daily correlations at some sites (0.20<R<0.79, Supplementary Fig. 13) and no correlations (R<0.10) at an hourly resolution except at two sites (Chicken Reef and Martin Reef). Therefore, we excluded temperature metrics based on hourly measurements (e.g. daily thermal range) from our environmental dataset.

6. **Random forest ensemble learning**

Low correlated environmental predictors were used in the random forest approach (absolute Pearson coefficients <0.7, Supplementary Table. 9) as multicollinearity can introduce bias in evaluation of variables importance. When two metrics were highly correlated, the most ecologically relevant one (or conceptually simple) was kept based on literature of environmental drivers of heat tolerance. St Crispin Reef data were removed from ED50 models as previous analyses showed low confidence in heat tolerance estimates at this site due to temperature treatments not matching targets. Random forest models were implemented using the cforest function from *party* R package ^3^. This package uses conditional inference trees in which permutation-based significance tests are used to split nodes (instead of information measure maximization), ensuring unbiased variable selection even when variables have different cardinality and scale of measurement ^4^. Repeated 10-folds stratified (across sites) cross validation (70/30 split) was used to assess mean variables importance and model predictive accuracies. Random forest parameters were set as 1000 for the number of trees per forest (ntree) and 5 for the number of variables tried at each split (mtry) based on sensitivity analysis (Supplementary Fig. 14).

### B. Figures

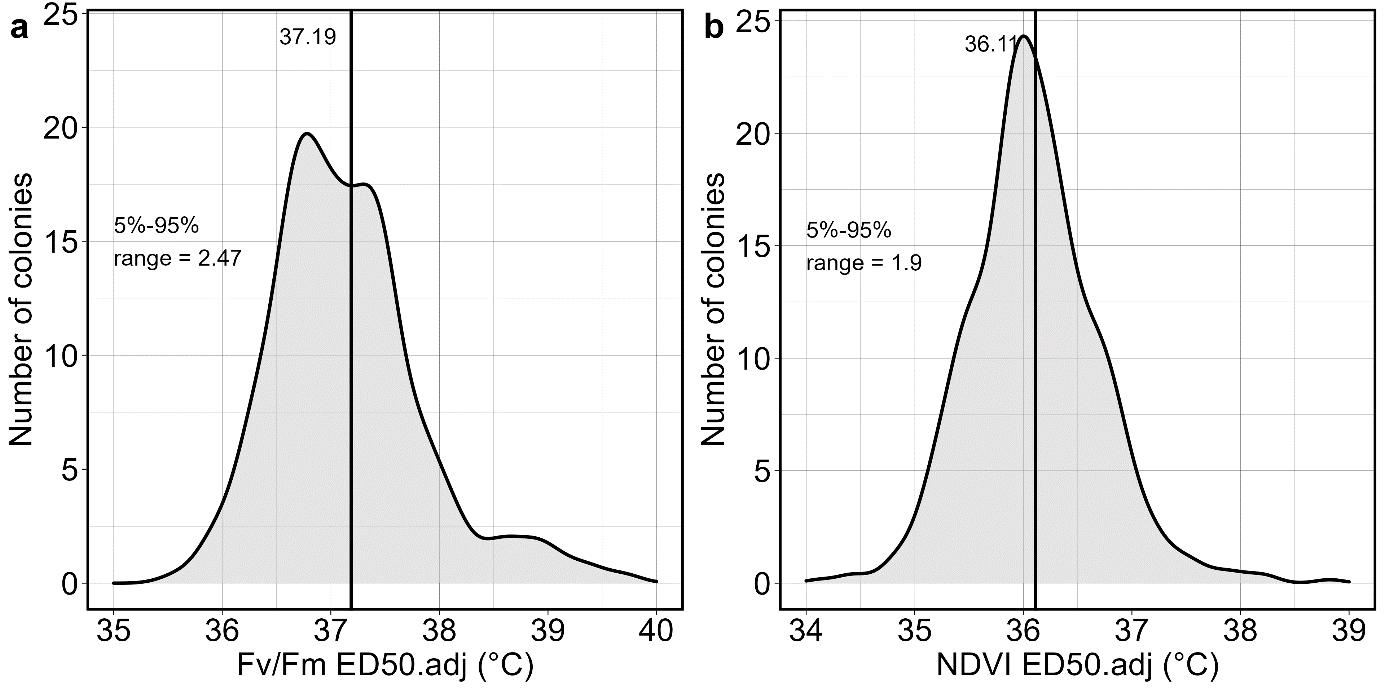

**Supplementary Figure 1.** Within-site variation in heat tolerance measured on maximum quantum yield (*Fv/Fm*) and chlorophyll a content (NDVI). **a.** Adjusted ED50 values from the total population (n=14 sites) by subtracting site effect from the total variation. Vertical line indicates the mean value of the adjusted distribution and 5%–95% quartiles range is shown. **b.** Same for NDVI.

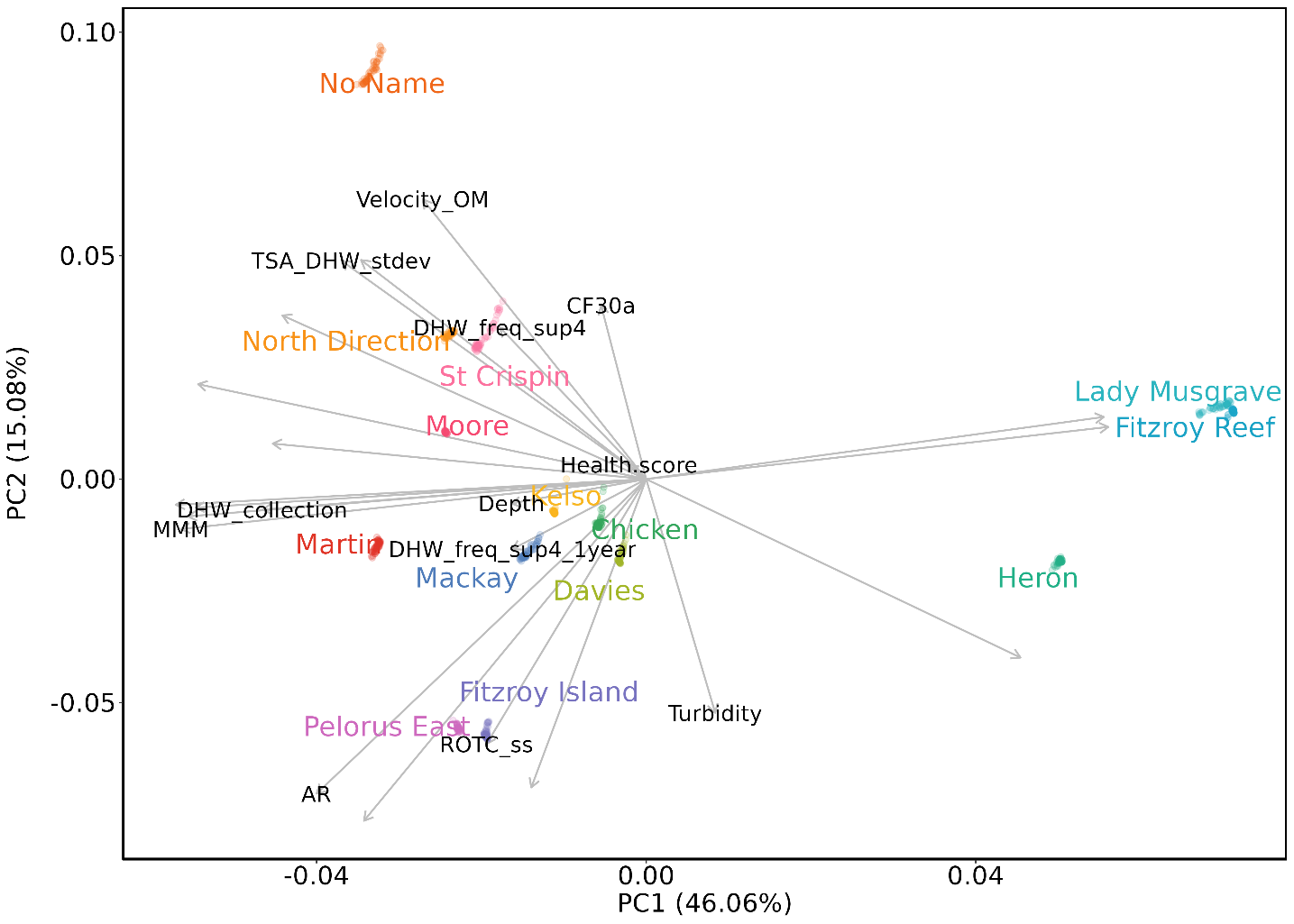

**Supplementary Figure 2.** Principal component analysis (PCA) conducted on 24 quantitative environmental predictors of *Acropora spathulata* heat tolerance metrics (*n* = 768 colonies sampled across the Great Barrier Reef, colored by their reef of origin). Loadings projected on the first two components are depicted with arrows, and labels are shown for moderately correlated predictors (|R| < 0.7) used in further analyses. CF30a = Cloud fraction anomaly for 30 days prior collection, DHW_collection = Degree Heating Weeks (DHW) at the day of collection, DHW_freq_sup4 = Frequency of DHW > 4, DHW_freq_sup4_1year = Frequency of DHW > 4 in the year prior collection, MMM = Maximum Monthly Mean (1985–1990+1993 climatology), ROTC_ss = Rate of temperature change in spring/summer, TSA_DHW_stdev = standard deviation of DHW. Numbers in parentheses represent the proportion of variance explained by each principal component.

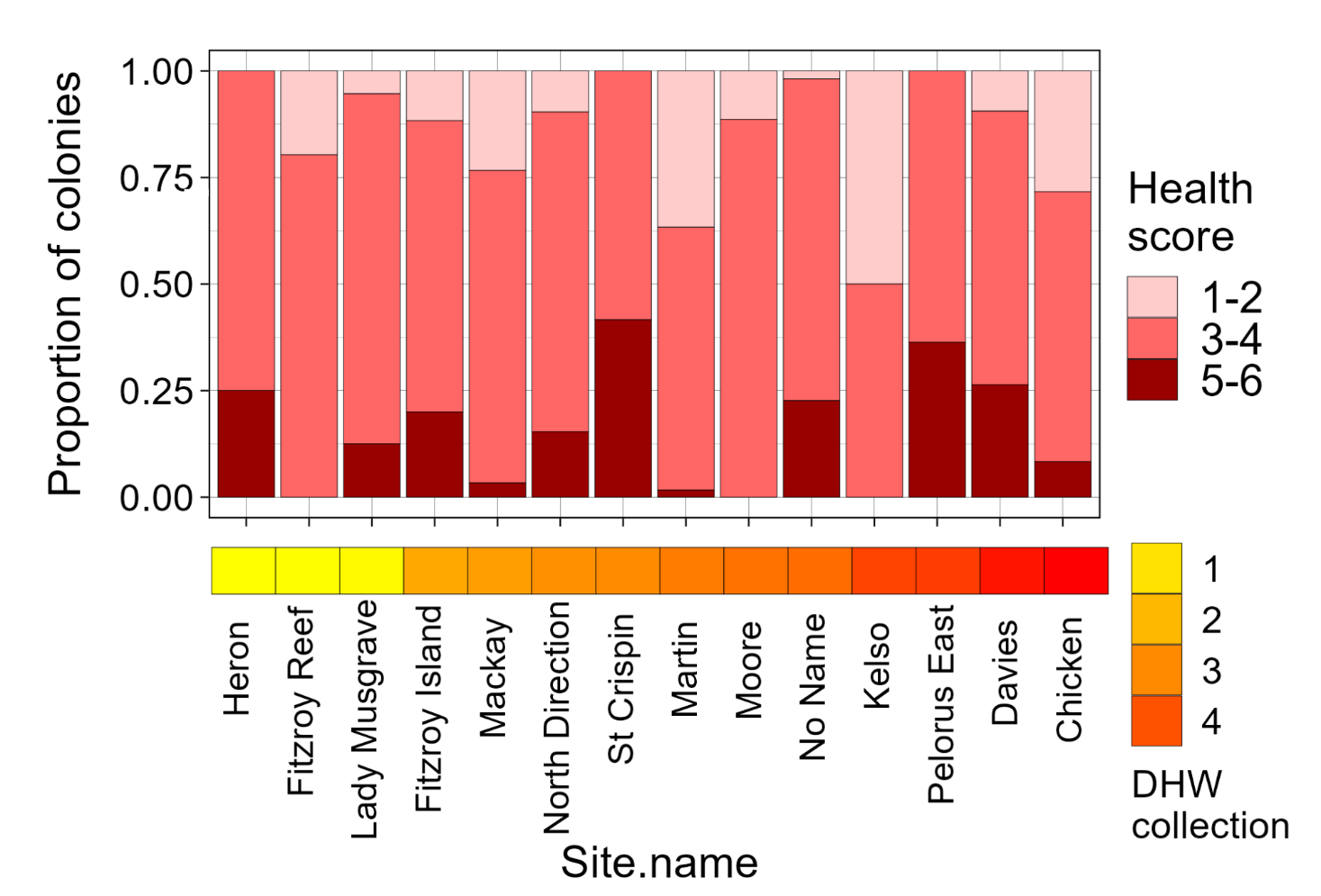

**Supplementary Figure 3.** Pigmentation of *A.spathulata* colonies prior to collection at the 14 sites were acute heat stress assay were conducted. The stacked bar chart shows the proportion of individual that were assigned a health score from 1=most bleached to 6=least bleached using the Coral Health Chart. Health score are grouped by 2 given the recording uncertainty. 1d heatmap shows ongoing Degree Heating Weeks (DHW) at the time colonies were collected, computed from NOAA Coral Watch v3.1 satellite product at 5km resolution.

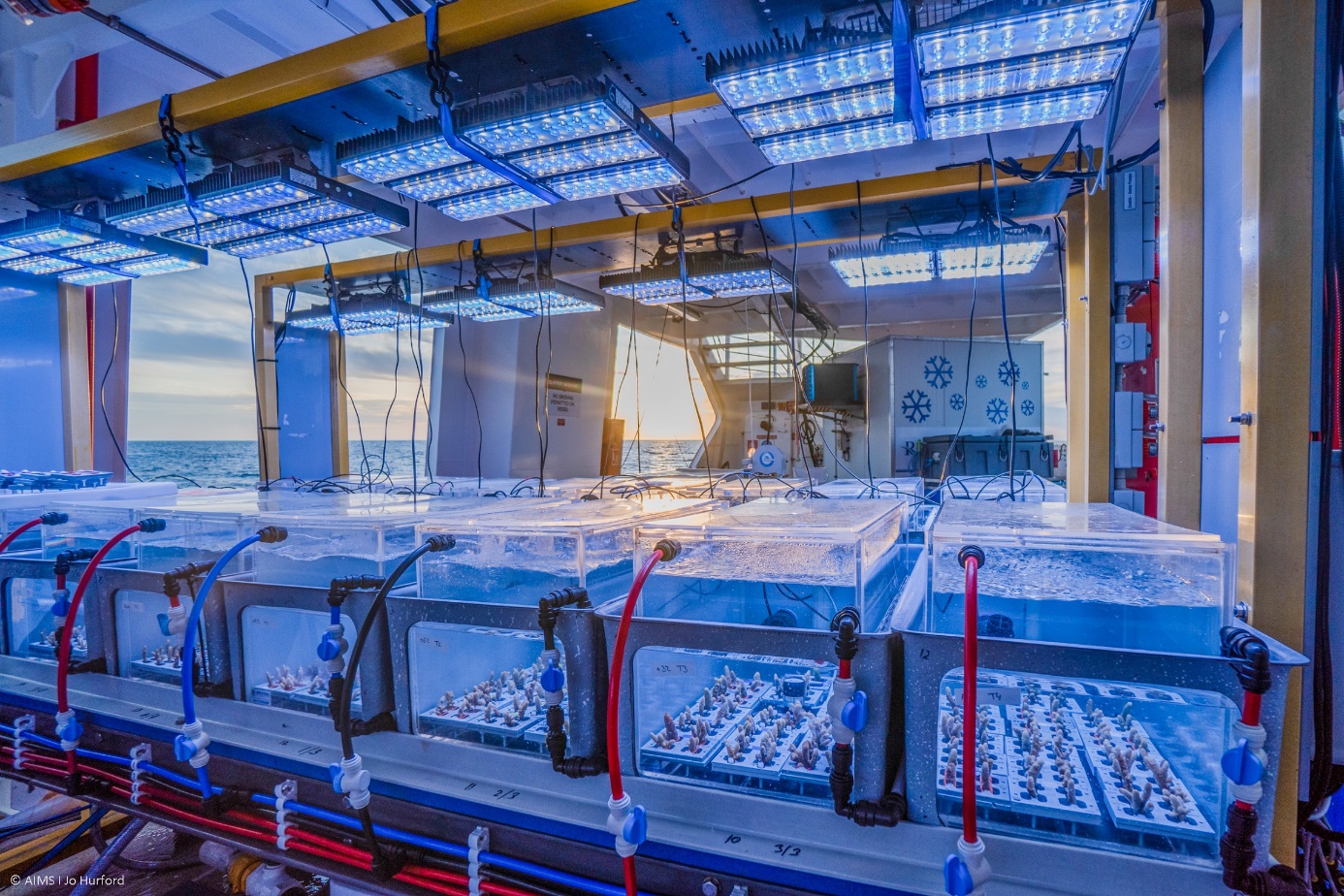

**Supplementary Figure 4.** Picture of the automated acute heat stress system (Seasim-in-a-box, SSIAB) to conduct automated, high-throughput acute heat stress experiments on corals directly in the field. The system is set up on research vessels which enables to conduct large scale phenotyping of coral populations. Southern Great Barrier Reef, March 2022. ©XXX Credit JoHurford 2022.

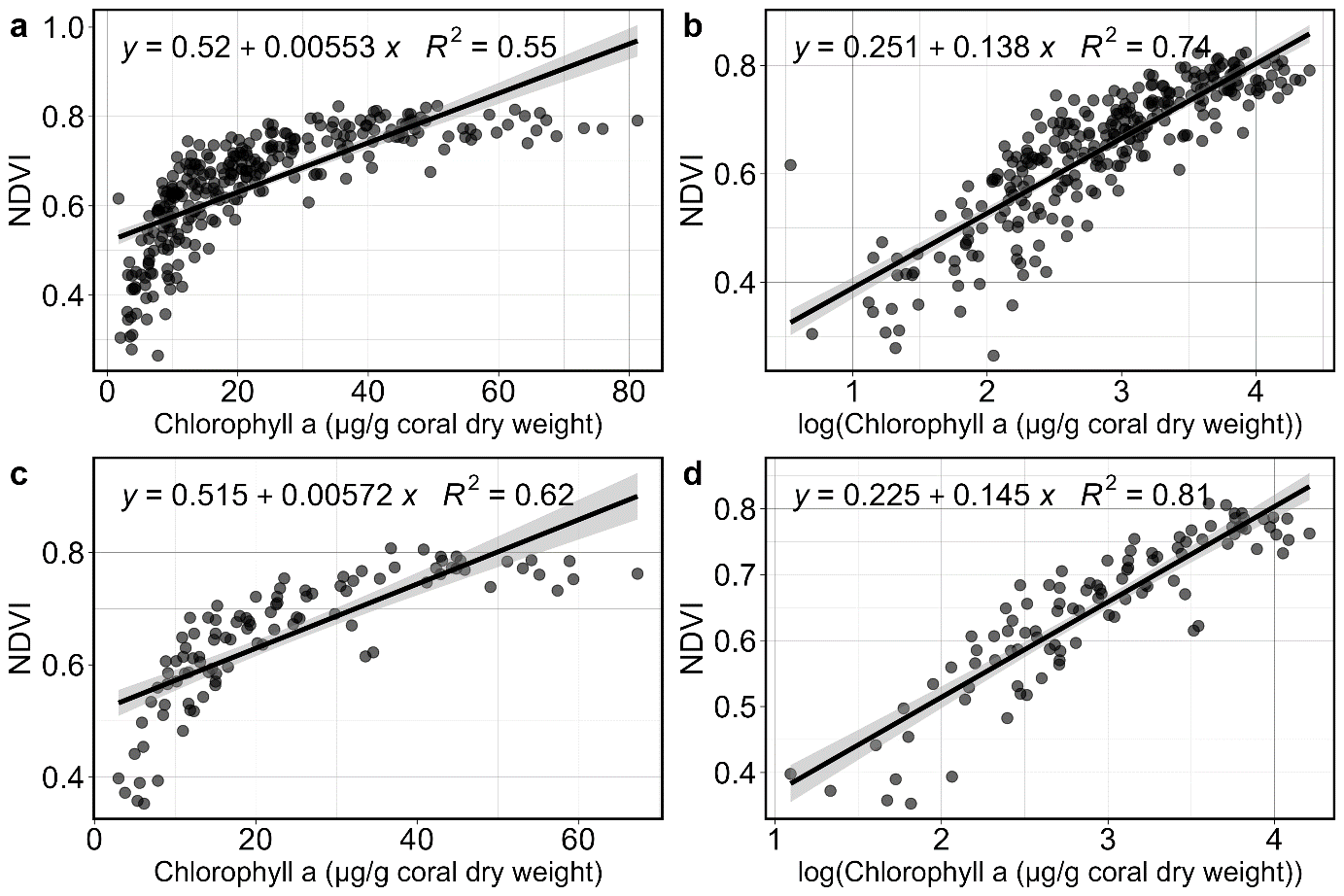

**Supplementary Figure 5.** Validation of chlorophyll-a quantification using hyperspectral imaging on coral fragments at the end of the heat stress experiment. Correlation between **a.** chlorophyll a and **b**. log transformed chlorophyll a and the normalized difference vegetation index (NDVI) for a subset of *A.spathulata* samples (n=285). Each point represents a single coral fragment originating from Chicken Reef or Pelorus Island and MMM or MMM+6°C. The linear equation and coefficient of determination (*R*^2^) are presented. **c.** and **d.** panels show the same correlation when values are averaged by colony and treatment (n=3 fragment/colony/treatment).

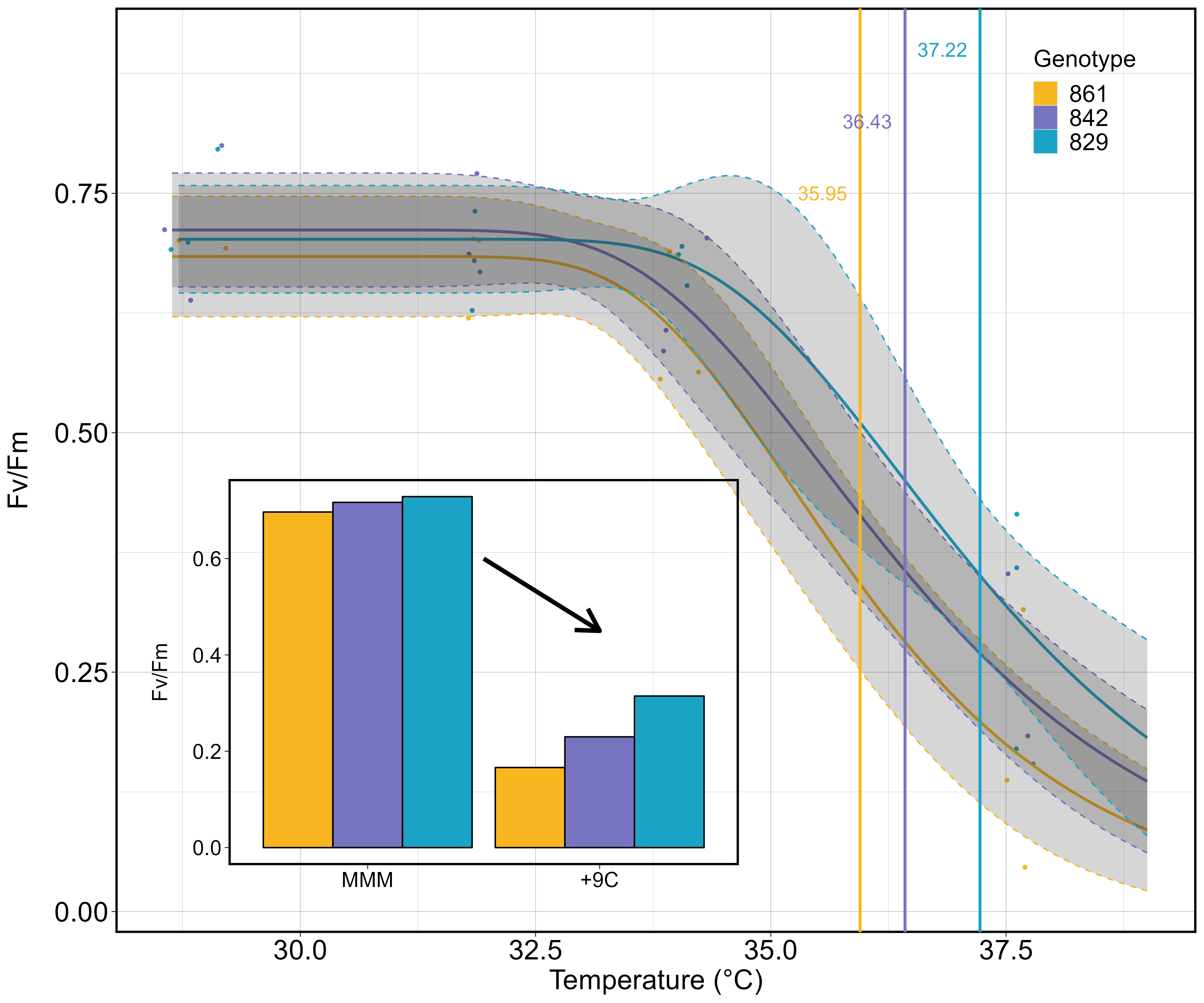

**Supplementary Figure 6.** Principle of heat tolerance measurements on *Acropora spathulata* across the Great Barrier Reef with the example of maximum photosynthetic efficiency of microalgal symbionts (*Fv/Fm*). The main panel depicts how individual *Fv/Fm* measurements from replicate fragments across four temperature treatments are used to compute Effective dose 50 (ED50): the absolute temperature eliciting a 50% drop in *Fv/Fm*. ED50s are indicated by vertical bars. The inset shows the principle of performance retention under extreme heat as the ratio of *Fv/Fm* in the +9 °C to the MMM treatment.

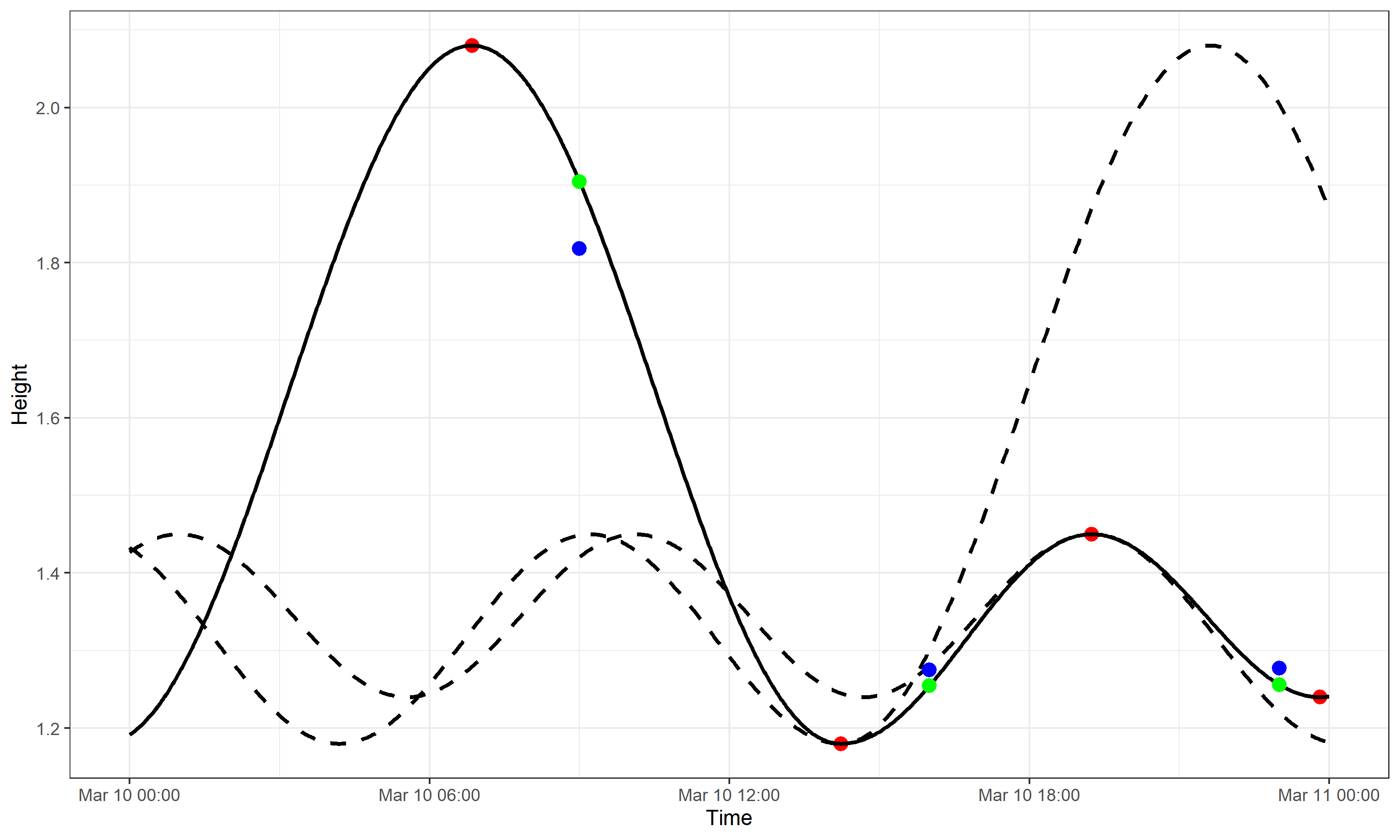

**Supplementary Figure 7.** Standardization of colonies depth to Lowest Astronomical Tide (LAT). For each individual colony, tide level at the time of collection was estimated from reef-level daily high and low tides (red dots) predicted by the Bureau of Meteorology, Australia^5^ and accessed at <https://tides.willyweather.com.au/>. The solid line displays the modeled tide from the four daily measurements. Green dots show the improvement of tide predictions using a sinusoid function rather than linear interpolation (blue dots). The standardized depth was computed as the colony depth measured from dive computers minus the tide level at time of collection, standardized to LAT. Negative depths indicate colonies above the LAT.

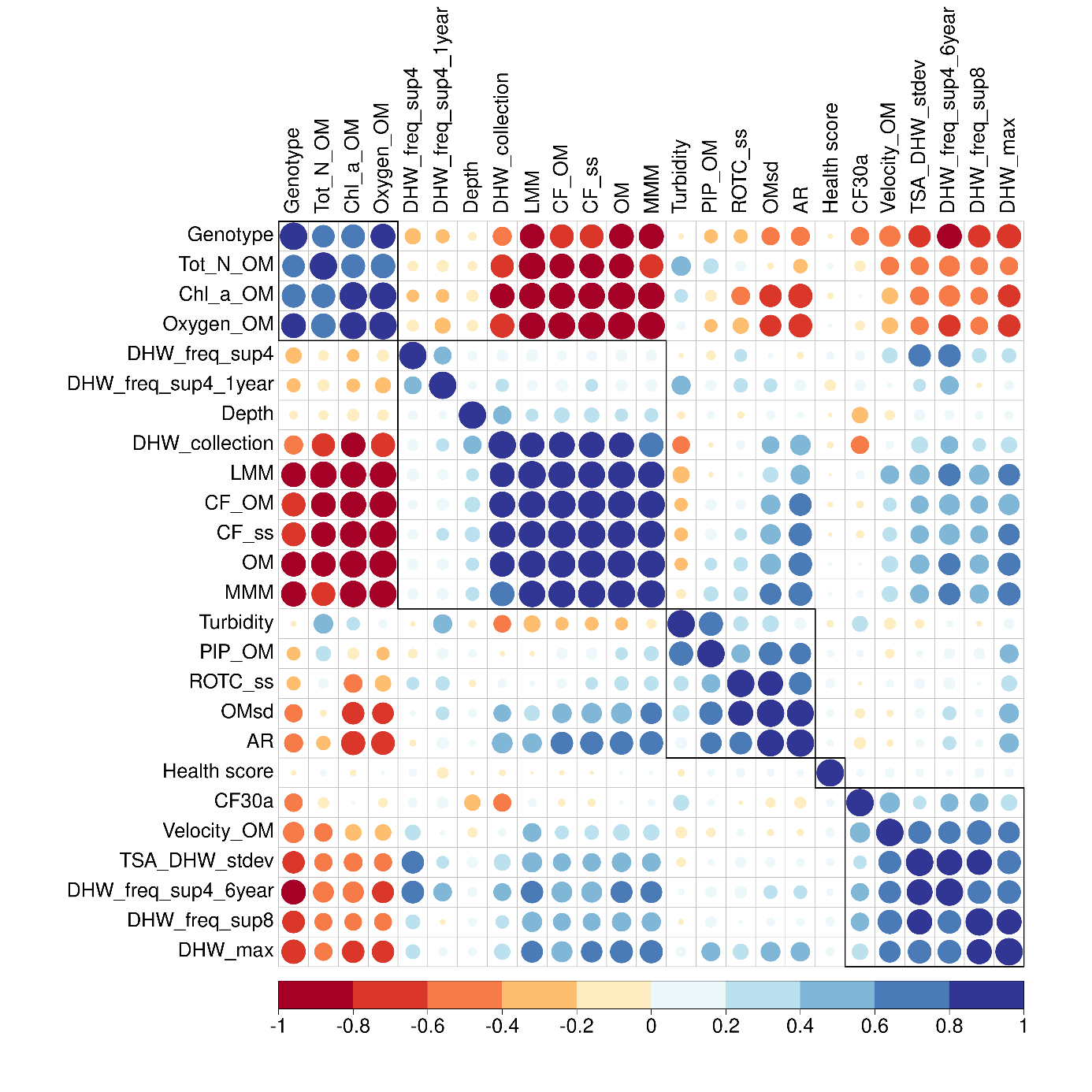

**Supplementary Figure 8.** Pairwise Pearson’s correlation coefficients between the 24 quantitative environmental predictors preliminarily extracted from various sources to assess environmental conditions experienced by *A.spathulata* colonies. Fourteen predictors were discarded from phenotype by environment models as they exhibited high collinearity with other covariates (absolute coefficient >0.7). For each pair of highly collinear variables, the most ecologically relevant one was kept in the model.

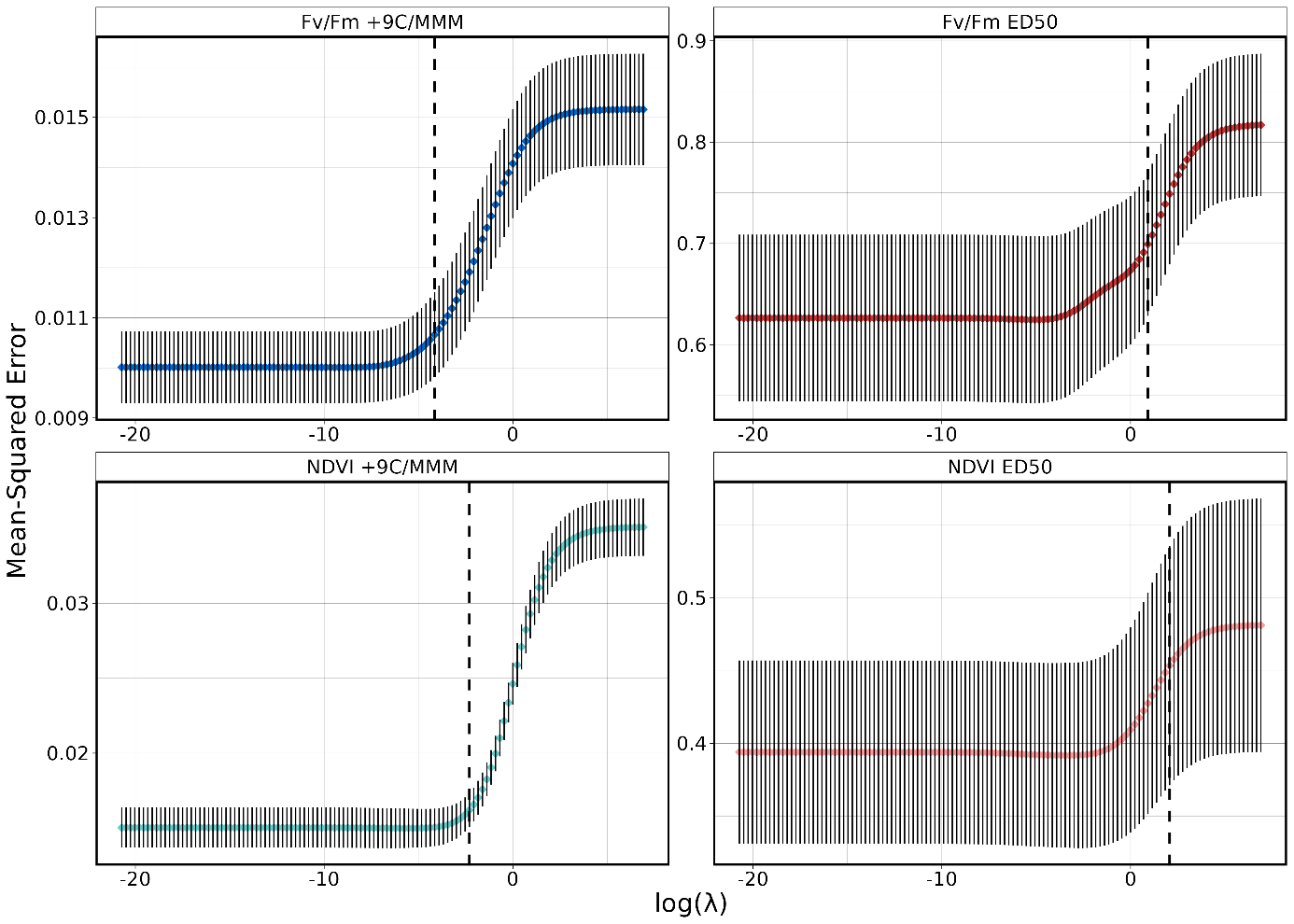

**Supplementary Figure 9.** Selection of tuning parameter *λ* of penalization coefficient for phenotypic traits ridge regression on the set of 31 environmental predictors. The optimal λ is chosen separately for each trait through *k*-fold cross-validation. For each λ value, the average mean squared error (colored dot) and standard deviation (horizontal bars) of ridge regression are computed on the test fold. The vertical dashed line shows the λ.1se parameter retained as the optimal λ, corresponding to the maximum lambda value with MSE within one standard error from min MSE (more regularization than λ.min).

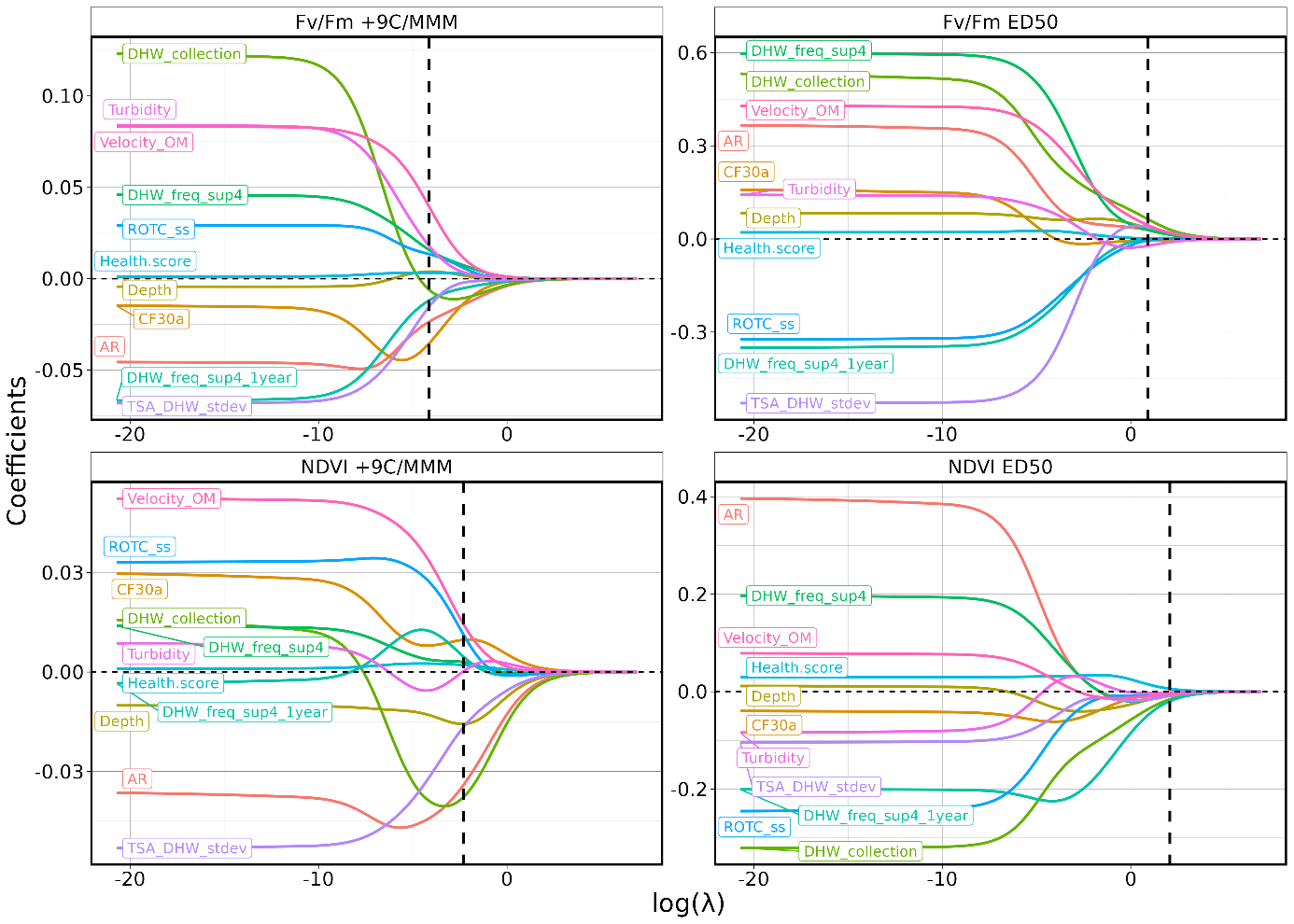

**Supplementary Figure 10.** Regularization of environmental predictors coefficients across increasing levels of penalization. Each colored line corresponds the value of a predictor coefficient for different values of tuning parameter *λ*. Variance in predictor estimates decreases as *λ* increases and bias increases as *λ* increases. The vertical dashed line shows the value of *λ*.1se obtained through cross-validation as providing the best bias-variance trade-off.

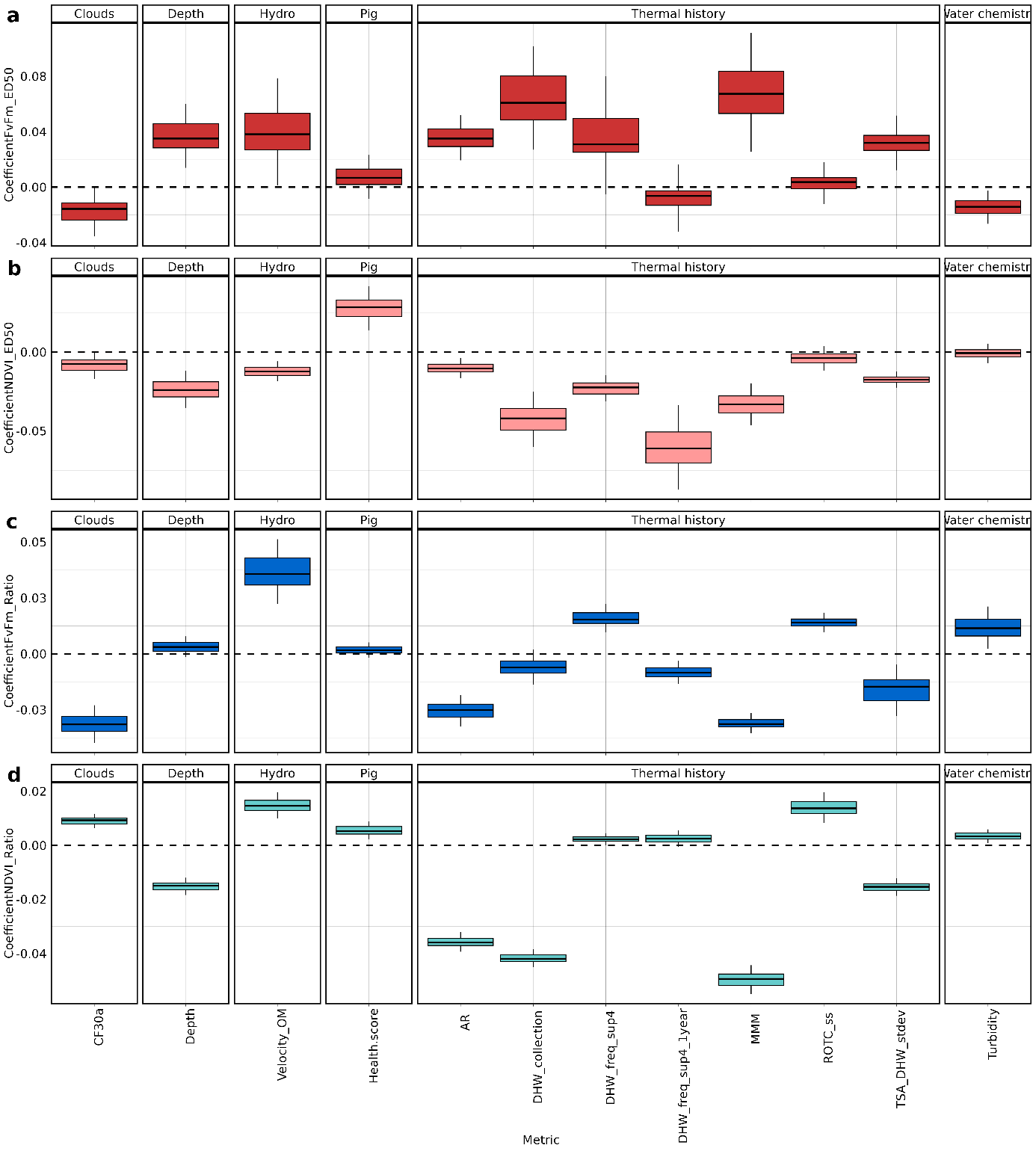

**Supplementary Figure 11.** Details of ridge regression predictors coefficients for **a.** *F_v_/F_m_* ED50, **b.** NDVI ED50, **c** *F_v_/F_m_* +9°C/MMM, **d.** NDVI +9°C/MMM . Estimates are grouped by types of environmental predictors and their variance assessed through bootstrap resampling (repeated 100-fold stratified cross validation). Boxplots display first quartile, median and third quartile for each predictor across the 100 folds. Pig=Pigmentation, Hydro=Hydrodynamics.

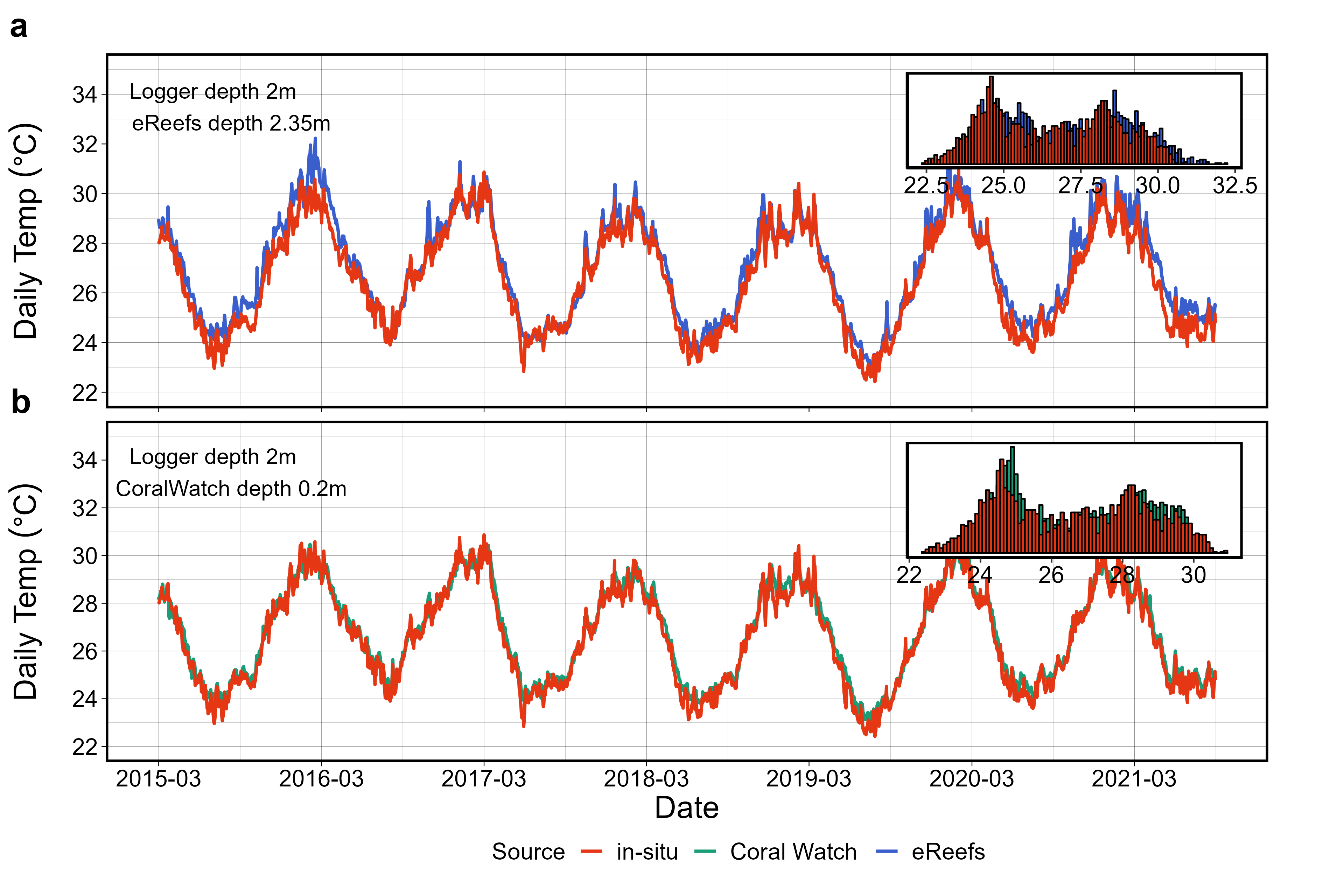

**Supplementary Figure 12.** Comparison of temperature records from various sources (in-situ loggers, satellite, numerical model) to characterize thermal history of the 14 collection sites. A representative example of temperature time-series is shown at Mackay Reef over the period 2015-2021. **a**. Daily *in-situ* logger averages (red) are plotted against eReefs temperatures (blue) extracted from GBR1 Hydro model at 1km resolution. **b**. Same against sea surface temperatures from Coral Watch v3.1 (green) at a 5km resolution. Histograms in the insets show the distribution of daily average temperatures for the various sources.

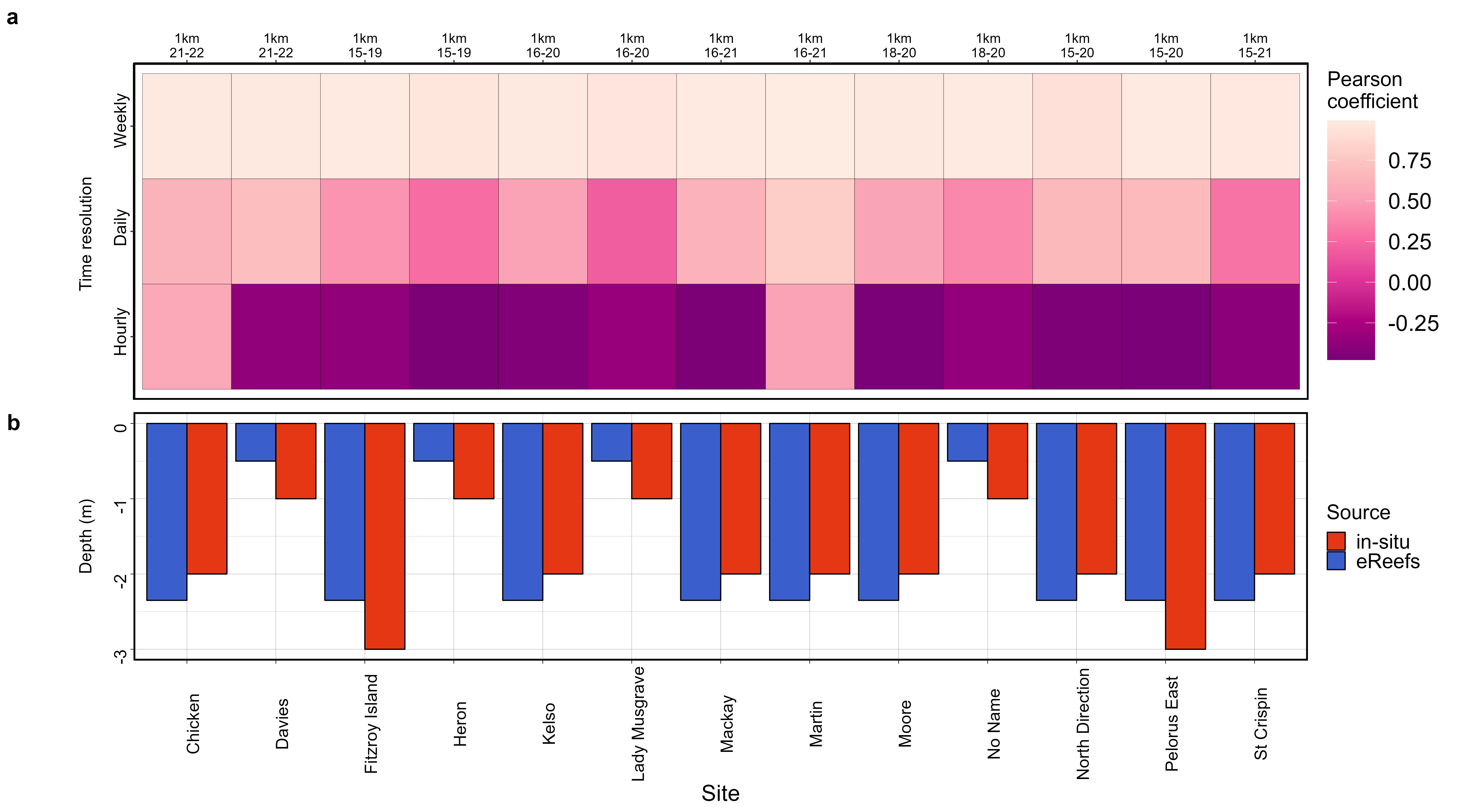

**Supplementary Figure 13.** Correlation between temperatures recorded by *in-situ* loggers and predicted by the eReefs model across time resolutions. **a.** Heatmap of Pearson’s correlation coefficients computed from time-series at hourly, daily and weekly resolutions for the 13 sites were *in-situ* data was available. To compute correlations, daily and hourly temperatures were adjusted to remove low frequency variations by subtracting weekly and daily averages respectively. Above the heatmap is indicated the resolution of the eReefs model and time period used for the comparison. All in-situ loggers had a minimum of one year of data. **b**. Depth of temperatures measurements from *in-situ* loggers and predictions from eReefs at each site. For *in-situ* loggers where information was not available, depth was arbitrarily set to 2m.

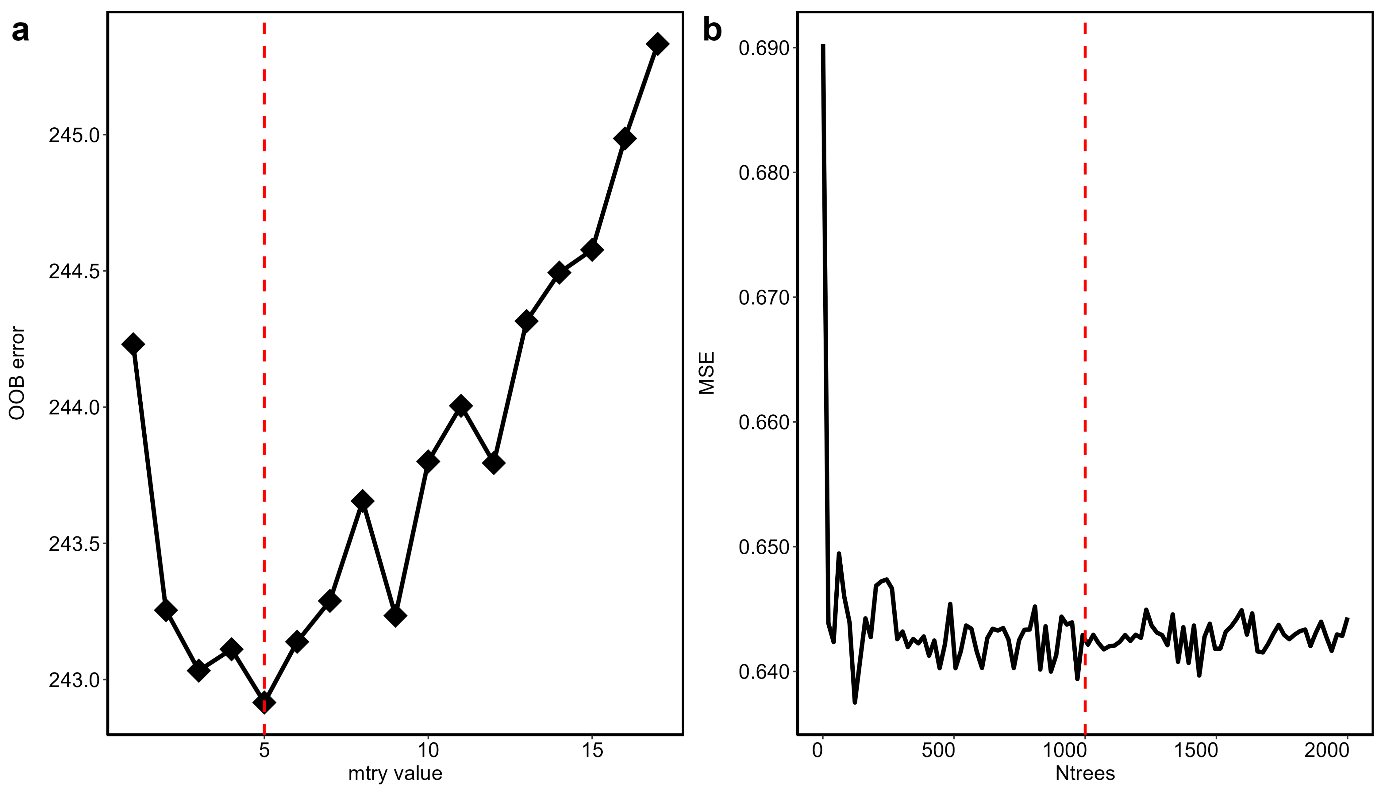

**Supplementary Figure 14.** Parameter tuning for conditional inference random forest*.* A representative examples of model sensibility to parameters is given for *Fv/Fm* ED50. **a.** Variation of out of bag sample average mean squared error (n=10 independent splits) with number of variables randomly tried at each node of the conditional inference tree (*mtry* parameter). The red line indicates the value of *mtry* parameter for which the model has the best predictive accuracy. **b**. Variation of test set mean squared error with the number of trees grown for each forest (*ntree* parameter). The red line indicates the value of *ntree* parameter that was used to train the model.
